# Control of Arabidopsis shoot stem cell homeostasis by two antagonistic CLE peptide signalling pathways

**DOI:** 10.1101/2021.06.14.448384

**Authors:** Jenia Schlegel, Grégoire Denay, Karine Gustavo Pinto, Yvonne Stahl, Julia Schmid, Patrick Blümke, Rüdiger Simon

## Abstract

Stem cell homeostasis in plant shoot meristems requires tight coordiantion between stem cell proliferation and cell differentiation. In Arabidopsis, stem cells express the secreted dodecapeptide CLAVATA3 (CLV3), which signals through the leucine-rich repeat (LRR)–receptor kinase CLAVATA1 (CLV1) and related CLV1-family members to downregulate expression of the homeodomain transcription factor *WUSCHEL* (*WUS*). WUS protein moves from cells below the stem cell domain to the meristem tip and promotes stem cell identity, together with *CLV3* expression, generating a negative feedback loop. How stem cell activity in the meristem centre is coordinated with organ initiation and cell differentiation at the periphery is unknown.

We show here that the *CLE40* gene, encoding a secreted peptide closely related to CLV3, is expressed in the SAM in differentiating cells in a pattern complementary to that of *CLV3*. *CLE40* promotes *WUS* expression via BAM1, a CLV1-family receptor, and *CLE40* expression is in turn repressed in a *WUS*-dependent manner. Together, *CLE40-BAM1-WUS* establish a second negative feedback loop. We propose that stem cell homeostasis is achieved through two intertwined pathways that adjust WUS activity and incorporate information on the size of the stem cell domain, via *CLV3-CLV1*, and on cell differentiation via *CLE40-BAM1*.

## Introduction

In angiosperms, the stem cell domain in shoot meristem is controlled by the directional interplay of two adjacent groups of cells. These are the central zone (CZ) at the tip of the dome-shaped meristem, comprising slowly dividing stem cells, and the underlying cells of the organising centre (OC). Upon stem cell division, daughter cells are displaced laterally into the peripheral zone (PZ), where they can enter differentiation pathways (Fletcher et al., 1999; Hall & Watt, 1989; Reddy et al., 2004; Schnablová et al., 2020; Stahl & Simon, 2005; Steeves & Sussex, 1989). Cells in the OC express the homeodomain transcription factor WUSCHEL (WUS), which moves through plasmodesmata to CZ cells to maintain stem cell fate and promote expression of the secreted signalling peptide CLAVATA3 (CLV3) (Brand et al., 2000; Daum et al., 2014; Müller et al., 2006; Schoof et al., 2000; Yadav et al., 2011). Perception of CLV3 by plasma-membrane localised receptors in the OC cells triggers a signal transduction cascade and downregulates WUS activity, thus establishing a negative feedback loop (Mayer et al., 1998; Ogawa et al., 2008; Yadav et al., 2011). Mutants of *CLV3* or its receptors (see below) fail to confine *WUS* expression and cause stem cell proliferation, while W*US* mutants cannot maintain an active stem cell population (Brand et al., 2002; Clark et al., 1993, 1995; Endrizzi et al., 1996; Laux et al., 1996; Schoof et al., 2000). WUS function in the OC is negative regulated by HAM transcription factors, and only WUS protein that moves upwards to the stem cell zone, which lacks *HAM* expression, can activate *CLV3* expression (Han et al., 2020; Zhou et al., 2018). The *CLV3-WUS* interaction can serve to maintain the relative sizes of the CZ and OC, and thereby meristem growth along the apical-basal axis. However, cell loss from the PZ due to production of lateral organs requires a compensatory size increase of the stem cell domain.

The CLV3 signalling pathway, which acts along the apical-basal axis of the meristem, has been widely studied in several plant species and shown to be crucial for stem cell homeostasis in shoot and floral meristems (Somssich et al., 2016). The CLV3 peptide is perceived by a leucin-rich-repeat (LRR) receptor kinase, CLAVATA1 (CLV1), which interacts with coreceptors of the CLAVATA3 INSENSITIVE RECEPTOR KINASES (CIK) 1-4 family (Clark et al., 1997; Cui et al., 2018). CLV1 activation involves autophoshorylation, interaction with membrane-associated and cytosolic kinases and phosphatases (Blümke et al., 2021; Defalco et al., 2021). Furthermore, heterotrimeric G-proteins and MAPKs have been implicated in this signal transduction cascade in maize and Arabidopsis (Betsuyaku et al., 2011; Bommert et al., 2013; Ishida et al., 2014; Lee et al., 2019). Besides CLV1, several other receptors contribute to WUS regulation, among them RECEPTOR-LIKE PROTEIN KINASE2 (RPK2), the CLAVATA2-CORYNE heteromer (CLV2-CRN) and BARELY ANY MERISTEM1-3 (BAM1-3) (Bleckmann et al., 2010; DeYoung & Clark, 2008; Hord et al., 2006; Jeong et al., 1999; Kinoshita et al., 2010; Müller et al., 2008). The BAM receptors share high sequence similarity with CLV1, and perform diverse functions throughout plant development. Double mutants of *BAM1* and *BAM2* maintain smaller shoot and floral meristems, thus displaying the opposite phenotype to mutants of CLV1 (DeYoung et al., 2006; DeYoung & Clark, 2008; Hord et al., 2006). Interestingly, ectopic expression experiments showed that CLV1 and BAM1 can perform similar functions in stem cell control (Nimchuk et al., 2015). In addition, one study showed that CLV3 could interact with CLV1 and BAM1 in cell extracts (Shinohara & Matsubayashi, 2015), although another *in vitro* study did not detect *BAM1-CLV3* interaction at physiological levels of CLV3 (Crook et al., 2020). Furthermore, CLV1 was shown to act as a negative regulator of *BAM1* expression, which was interpreted as a genetic buffering system, whereby a loss of CLV1 is compensated by upregulation of BAM1 in the meristem centre (Nimchuk, 2017; Nimchuk et al., 2015). Comparable genetic compensation models for CLE peptide signalling in stem cell homeostasis were established for other species, such as tomato and maize (Rodriguez-Leal et al., 2019).

Maintaining the overall architecture of the shoot apical meristem during the entire life cycle of the plant requires replenishment of differentiating stem cell descendants in the PZ, indicating that cell division rates and cell fate changes in both regions are closely connected (Stahl & Simon, 2005). Overall meristem size is restricted by the ERECTA-family signalling pathway, which is activated by EPIDERMAL PATTERNING FACTOR (EPF)-LIKE (EPFL) ligands from the meristem periphery and confines both *CLV3* and *WUS* expression (Mandel et al., 2014; Shpak, 2013; Shpak et al., 2004; Torii et al., 1996; Zhang et al., 2021). In the land plant lineage, the shoot meristems of bryophytes such as the moss *Physcomitrium patens* appear less complex than those of angiosperms, and carry only a single apical stem cell which ensures organ initiation by continuous asymmetric cell divisions (de Keijzer et al., 2021; Harrison et al., 2009). Broadly expressed CLE peptides were here found to restrict stem cell identity, and act in division plane control (Whitewoods et al., 2018). Proliferation of the apical notch cell in the liverwort *Marchantia polymorpha* is promoted by MpCLE2 peptide which acts from outside the stem cell domain via the receptor MpCLV1, while cell proliferation is confined by MpCLE1 peptide through a different receptor (Hata & Kyozuka, 2021; Hirakawa et al., 2019, 2020; Takahashi et al., 2021). Thus, antagonistic control of stem cell activities through diverse CLE peptides is conserved between distantly related land plants. In the grasses, several CLEs were found to control the stem cell domain. In maize, *ZmCLE7* is expressed from the meristem tip, while *ZmFCP1* is expressed in the meristem periphery and its centre. Both peptides restrict stem cell fate via independent receptor signalling pathways (Liu et al., 2021; Rodriguez-Leal et al., 2019). In rice, overexpression of the CLE peptides OsFCP1 and OsFCP2 downregulates the homeobox gene *OSH1* and arrests meristem function (Ohmori et al., 2013; Suzaki et al., 2008). Common for rice and maize, CLE peptide signalling restricts stem cell activities in the shoot meristem, but a stem cell promoting pathway were not been identified so far.

Importantly, how stem cell activities in the CZ and OC are coordinated to regulate organ initiation and cell differentiation in the PZ, which is crucial to maintain an active meristem, is not yet known. In maize, the CLV3-related peptide ZmFCP1 was suggested to be expressed in primordia, and convey a repressive signal on the stem cell domain (Je et al., 2016). In Arabidopsis, the most closely related peptide to CLV3 is CLE40, which was shown to act in the root meristem to restrict columella stem cell fate and regulate the expression of the *WUS* paralog *WOX5* (Berckmans et al., 2020; Hobe et al., 2003; Pallakies & Simon, 2014; Stahl et al., 2013; Stahl & Simon, 2010). Functions of CLE40 in the SAM have not previously been described. Overexpression of *CLE40* causes shoot stem cell termination, while *CLE40* expression from the *CLV3* promoter fully complements the shoot and floral meristem defects of *clv3* mutants (Hobe et al., 2003). We therefore hypothesized that *CLE40* could act in a *CLV3*-related pathway in shoot stem cell control.

Here, we show that the expression level of *WUS* in the OC is subject to feedback regulation from the PZ, which is mediated by the secreted peptide CLE40. In the shoot meristem, *CLE40* is expressed in a complementary pattern to *CLV3*, and excluded from the CZ and OC. In *cle40* loss of function mutants, *WUS* expression is reduced, and shoot meristems remain small and flat, indicating that *CLE40* signalling is required to maintain *WUS* expression in the OC. Ectopic expression of *WUS* represses *CLE40* expression, while in *wus* loss-of-function mutants *CLE40* is expressed in the meristem centre, indicating that CLE40, in contrast to CLV3, is subject to negative feedback regulation by WUS. CLE40 likely acts as an autocrine signal that is perceived by BAM1 in a domain flanking the OC.

Based on our findings, we propose a new model for the regulation of the stem cell domain in the shoot meristem in which signals and information from both, the CZ and the PZ are integrated through two interconnected negative feedback loops that sculpt the dome-shaped shoot meristems of angiosperms.

## Results

### CLE40 signalling promotes IFM growth from the peripheral zone

Previous studies showed that *CLE40* expression from the *CLV3* promoter can fully complement a *clv3-2* mutant, indicating that CLE40 can substitute CLV3 function in the shoot meristem to control stem cell homeostasis, if expressed from the stem cell domain. Furthermore, while all other *CLE* genes in Arabidopsis lack introns, the *CLE40* and *CLV3* genes carry two introns at very similar positions (Hobe et al., 2003). Phylogenetic analysis revealed that CLV3 and CLE40 locate in the same cluster together with CLV3 orthologues from rice, maize and tomato (Goad et al., 2017) (Fig. *1*A).

**Fig. 1:**
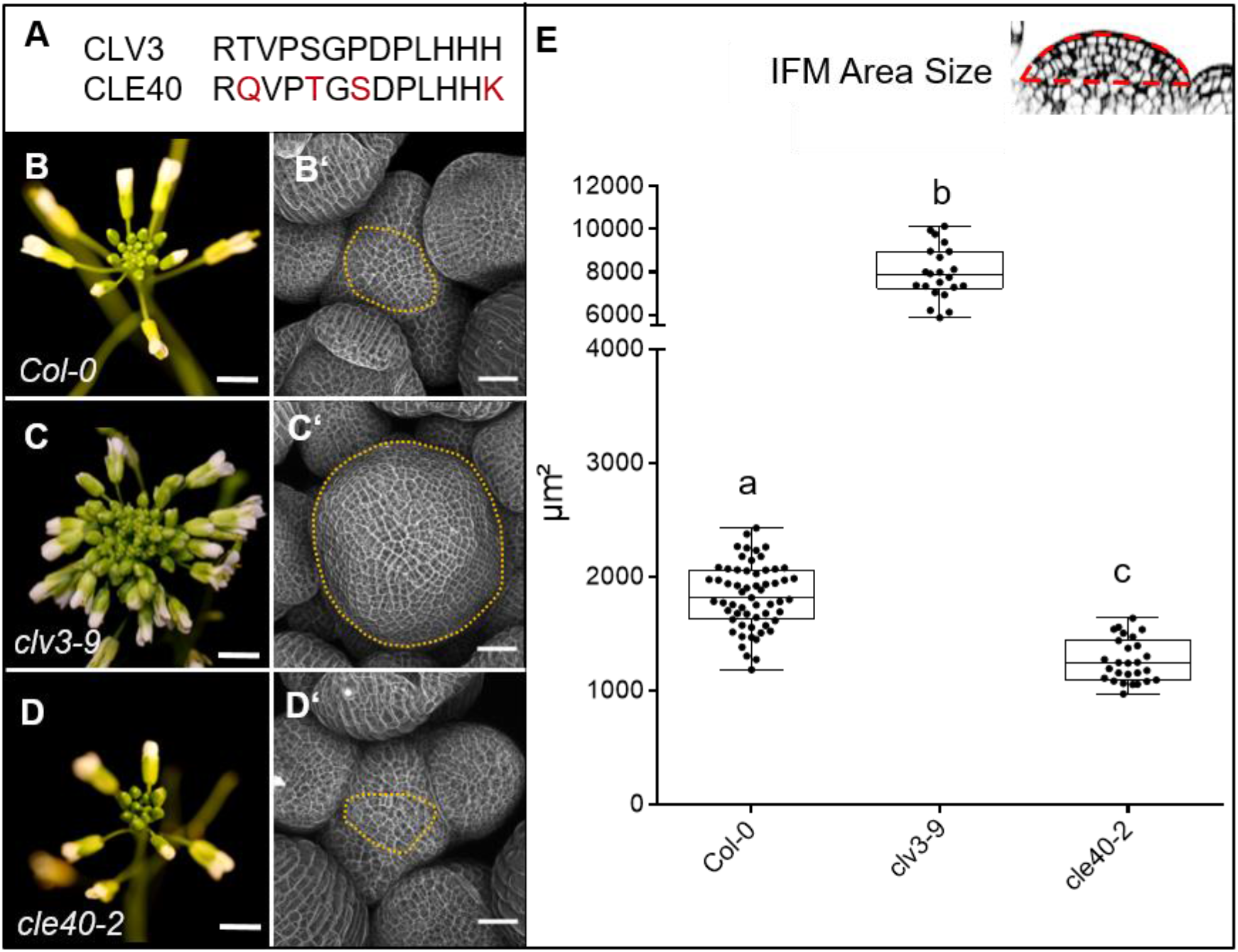
CLV3 and CLE40 exert opposite effects on meristem size. **(A)** The amino acid (AA) sequences of the mature CLV3 and CLE40 peptides differ in four AAs (differences marked in red). **(B)** *Col-0* inflorescence at 6 WAG with flowers. **(B’)** IFM at 6 WAG, maximum intensity projection (MIP) of a z-stack taken by confocal microscopy. **(C)** *clv3-9* inflorescence at 6 WAG **(Ć)** MIP of a *clv3-9* IFM at 6 WAG. **(D)** Inflorescence of *cle40-2* at 6 WAG **(D’)** MIP of a *cle40-2* IFM. **(E)** Box and whisker plot of IFM sizes of *Col-0* (N=59), *clv3-9* (N=22), and *cle40-2* (N=27) plants. Scale bars: 10mm (B, C, D), 50µm (B’, C’, D’), Statistical groups were assigned after calculating p-values by ANOVA and Turkey’s multiple comparison test (differential grouping from p ≤ 0.01). Yellow dotted lines in B’ to D’ enclose the IFM, red line in the inset meristem in E indicates the area that was used for the quantifications in E.

Mutations in *CLE40* were previously found to affect distal stem cell maintenance in the root meristem, revealing that a CLV3 related signalling pathway also operates in the root stem cell niche. To uncover a potential role of *CLE40* in shoot development, we analysed seedling and flower development, and inflorescence meristem (IFM) sizes of the wild type *Col-0*, and *clv3-9* and *cle40-2 loss-of-function* mutants. At 4 weeks after germination (WAG), leaves of *clv3-9* mutants remained shorter than those of *Col-0* or *cle40-2* (Fig1-SupplFig.1). After floral induction, the inflorescences of *clv3-9* mutants were compact with many more flowers than the wild type, while *cle40-2* mutant inflorescences appeared smaller than the control (Fig. *1*B-D). To first investigate effects on meristem development in detail, longitudinal optical sections through the inflorescence meristem (IFM) at 6 WAG were obtained by confocal microscopy and meristem areas were analysed (Fig. *1*B-E). In *clv3-9* mutants, meristem areas increased to approx. 450% of wild type (*Col-0*) levels, while shoot meristems from 4 independent *cle40* mutant alleles in a *Col-0* background (*cle40-2, cle40-cr1, cle40-cr2, cle40-cr3*) reached only up to 65% of wild type (Fig. *1*E, Fig1-SupplFig.2C) (Yamaguchi et al., 2017). Next, we used carpel number as a rough proxy for flower meristem (FM) size, which was 2±0.0 (N=290) in *Col-0* and *cle40-2* (N=290) but 3.7±0.4 (N=340) in *clv3-9* (Fig1-SupplFig.3). Hence, we concluded that CLE40 mainly promotes IFM growth, whereas CLV3 serves to restrict both IFM and FM sizes.

We next analysed the precise *CLE40* expression pattern using a transcriptional reporter line, *CLE40:Venus-H2B* (Wink, 2013). We first concentrated on the IFMs and FMs. *CLE40* is expressed in IFMs and in FMs, starting at P5 to P6 onwards (Fig. *2*A-C). We found stronger expression in the PZ than in the CZ, and no expression in young primordia. Using MorphoGraphX software, we extracted the fluorescence signal originating from the outermost cell layer (L1) of the IFM, and noted reduced *CLE40* expression in the CZ (Fig. *2*B). Optical longitudinal sections through the IFM showed that *CLE40* is not expressed in the CZ, and only occasionally in the OC region (Fig. *2*C). Expression of *CLE40* changed dynamically during development: expression was concentrated in the IFM, but lacking at sites of primordia initiation (P0 to P4/5, Fig. *2*C). In older primordia from P5/6 onwards, *CLE40* expression is detectable from the centre of the young FM and expands towards the FM periphery. In the FMs, *CLE40* is lacking in young sepal primordia (P6), but starts to be expressed on the adaxial sides of petals at P7 (Fig. *2*A, P1-P7).

**Fig. 2:**
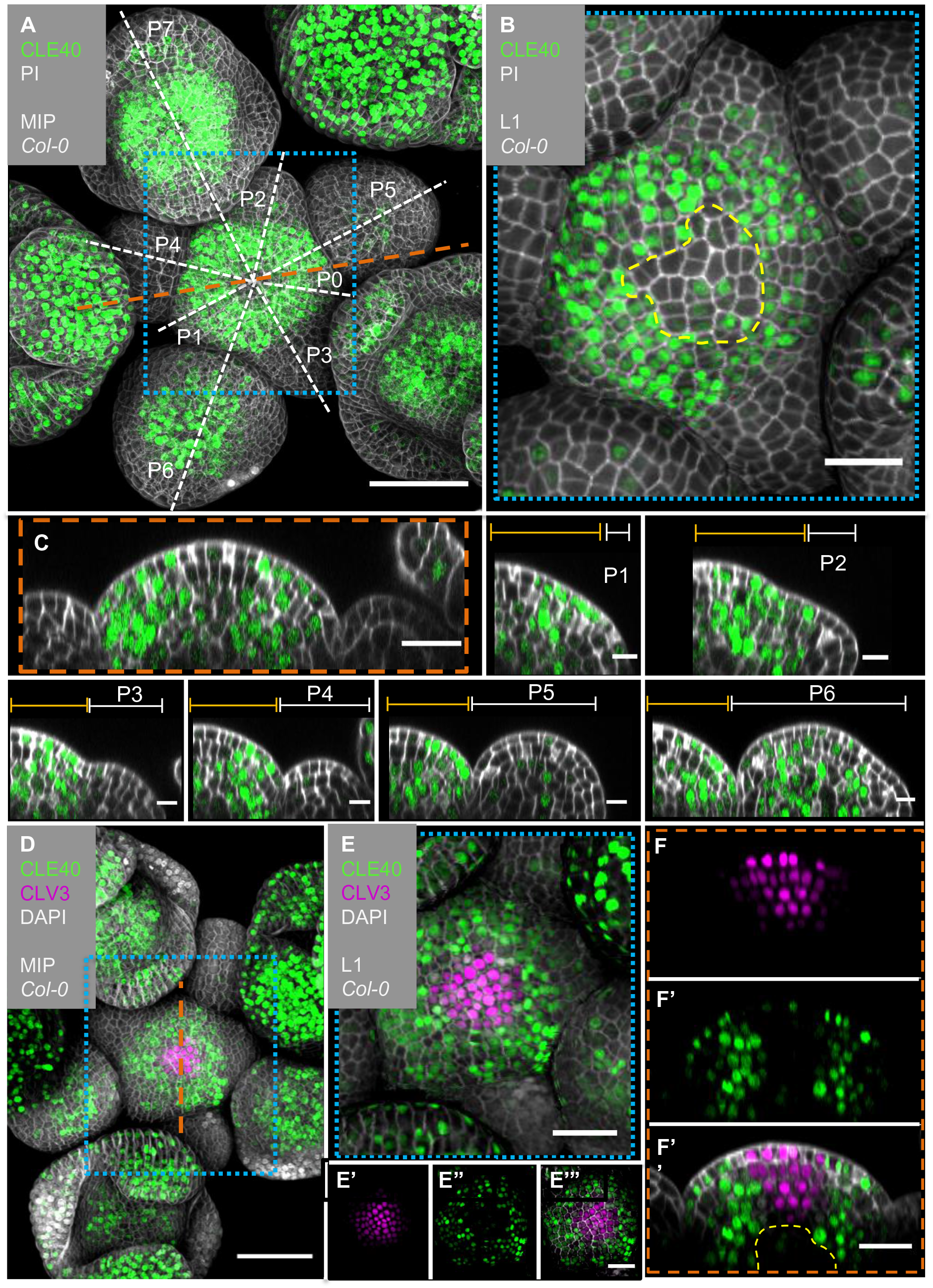
*CLE40* and *CLV3* show complementary expression patterns in the IFM. **(A)** MIP of an inflorescence at 5 WAG expressing the transcriptional reporter *CLE40:Venus-H2B//Col-0* showing *CLE40* expression in the IFM, older primordia and sepals (N=23). **(B)** The L1 projection shows high expression in the epidermis of the periphery of the IFM and only weak expression in the CZ. **(C)** Longitudinal section through the IFM shows expression of *CLE40* in the periphery, but lack of expression in the CZ. **(P1 –P6)** Longitudinal section through primordia show no *CLE40* expression in young primordia (P1-P4), but in the centre of older primordia (P5-P6). **(D)** The MIP of the double reporter line of *CLE40* and *CLV3* (*CLE40:Venus-H2B;CLV3:NLS-3xmCherry//Col-0*) shows *CLV3* expression in the CZ surrounded by *CLE40* expression in the periphery (N=12). **(E-E’’’)** The L1 projection shows *CLV3* **(E’)** expression in the centre of the IFM and *CLE40* **(E’’)** expression in a distinct complementary pattern in the periphery of the IFM. **(F)** The longitudinal section through the centre of the IFM shows *CLV3* expression in the CZ while *CLE40* **(F’)** is mostly expressed in the surrounding cells. **(F’’)** *CLE40* and *CLV3* are expressed in complementary patterns. Dashed blue lines indicate magnified areas, dashed white and orange lines indicate planes of optical sections, dashed yellow line in B marks CZ and in F’’ the OC. Scale bars: 50µm (A, D), 20µm (B, C, E, E’’’, F’’), 10µm (P0 to P6), MIP = Maximum intensity projection, PI = Propidium iodide, L1 = visualisation of layer 1 only, P1 to P7 = primordia at consecutive stages.

To compare the *CLE40* pattern with that of *CLV3*, we introgressed a *CLV3:NLS-3xmCherry* transcriptional reporter into the *CLE40:Venus-H2B* background. *CLV3* and *CLE40* are expressed in almost mutually exclusive domains of the IFM, with *CLV3* in the CZ surrounded by *CLE40* expressing cells (Fig. *2*D-F’’). In the deeper region of the IFM, where the OC is located, both *CLV3* and *CLE40* are not expressed (Fig. *2*F).

We noted that *CLE40* is downregulated where *WUS* is expressed, or where WUS protein localises, such as the OC and CZ. Furthermore, *CLE40* is also lacking in very early flower primordia and in incipient organs.

### *CLE40* expression is repressed by WUS activity

To further analyse the regulation of *CLE40* expression, we introduced the *CLE40* transcriptional reporter into the *clv3-9* mutant background (Fig. *3*A-B, Fig3-SupplFig.1). In *clv3-9* mutants, *WUS* is no longer repressed by the *CLV* signalling pathway, and the CZ of the meristem increases in size as described previously (Clark et al., 1995). In the *clv3-9* mutant meristems, both *CLV3* and *WUS* promoter activity is now found in an expanded domain (Fig3-SupplFig.1). *CLE40* is not expressed in the tip and centre of the IFM but is rather confined to the peripheral domain, where neither *CLV3* nor *WUS* are expressed (Fig. *3*B’, Fig3-SupplFig.1B’). To further explore the expression dynamics of *CLE40* in connection with regulation of stem cell fate and WUS, we misexpressed *WUS* from the *CLV3* promoter and introgressed it into plants carrying the *CLE40:Venus-H2B* construct. Since WUS activates the *CLV3* promoter, *CLV3:WUS* misexpression triggers a positive feedback loop. This results in a continuous enlargement of the CZ (Brand et al., 2002). Young seedlings carrying the *CLV3:WUS* transgene at 10 DAG displayed a drastically enlarged SAM, compared to wild type seedlings of the same age (Fig. *3*C-D’). Wild type seedlings at this stage express *CLE40* in older leaf primordia and in deeper regions of the vegetative SAM (Fig. *3*E-E’). The *CLV3:WUS* transgenic seedlings do not initiate lateral organs from the expanded meristem, and *CLE40* expression is confined to the cotyledons (Fig. *3*F-F’). *CLE40* is also lacking in the deeper regions of the vegetative SAM (Fig. *3*F’). Thus. we conclude that either WUS itself, or a WUS-dependent regulatory pathway represses *CLE40* gene expression.

**Fig. 3:**
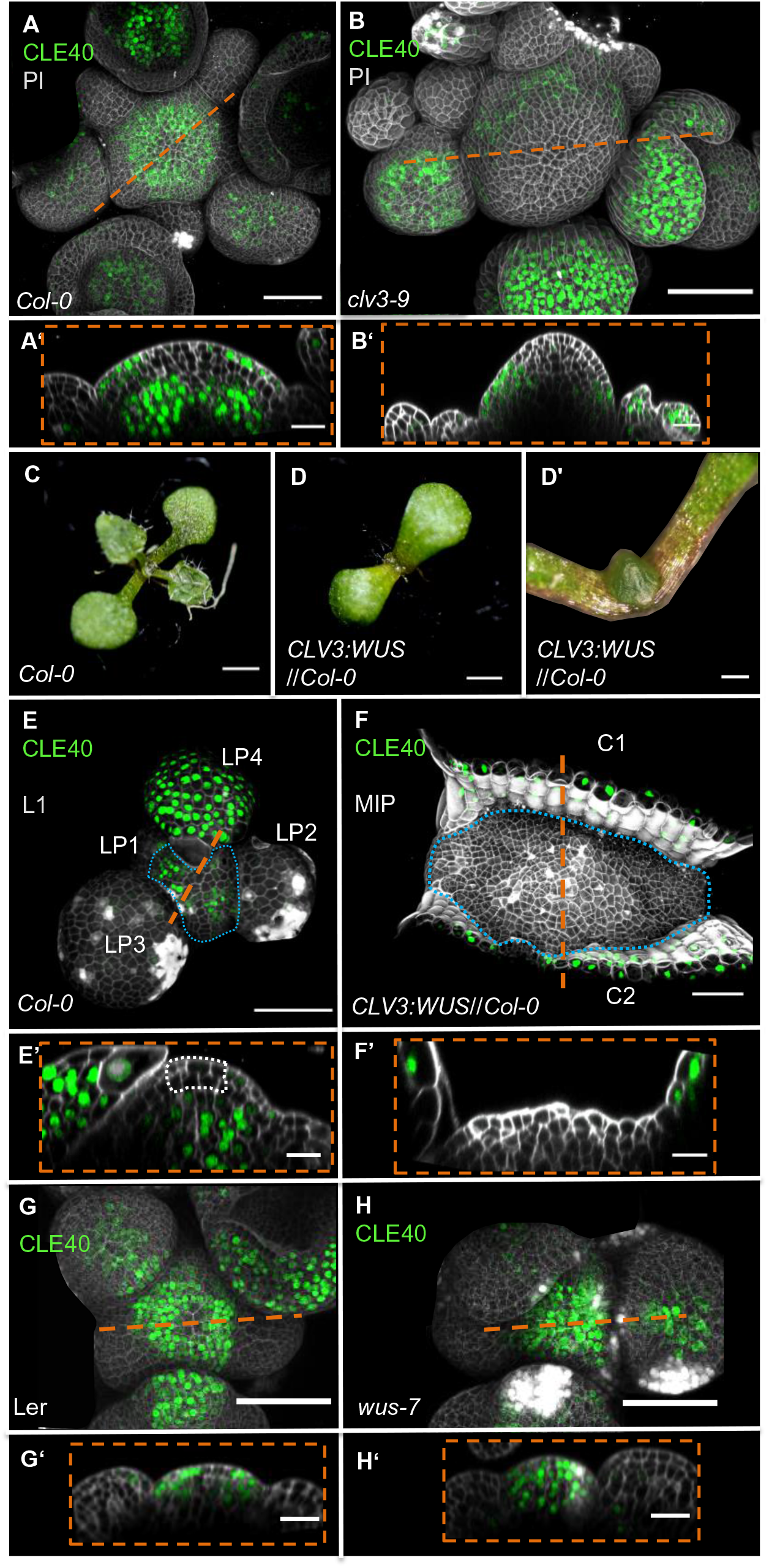
*WUS*-dependent repression of *CLE40* expression in the shoot meristem. **(A)** MIP of *CLE40* expression (*CLE40:Venus-H2B//Col-0*) at 5 WAG, **(A’)** Optical section through the centre of the IFM (indicated by orange line in (A)) reveals no *CLE40* expression in the CZ and in the centre of the meristem. Cells in the L2 layer also show less *CLE40* expression. High *CLE40* expression is found in the PZ (N=23). **(B)** MIP of *CLE40* expression in a *clv3-9* mutant (*CLE40:Venus-H2B//clv3-9*) shows expression only in the PZ of the meristem, in FMs and in sepals (N=6). **(B’)** Optical section through the IFM depicts no *CLE40* expression at the tip and the centre of the meristem. *CLE40* expression is only detected in cells at the flanks of the IFM and in sepals. **(C)** Arabidopsis seedling at 10 DAG. **(D)** Seedling expressing *WUS* from the *CLV3* promoter, 10 DAG. **(D’)** Magnification of seedling in (D). The meristem fasciates without forming flowers. **(E)** L1 projection, vegetative seedling with *CLE40* expression in the PZ and in leaf primordia starting from LP4, at 10 DAG (N=5). **(E’)** Optical section of (E) with *CLE40* expression primordia and rib meristem or periphery. **(F)** MIP of fasciated meristem as in (D). *CLE40* expression can only be found in the cotyledons (C1 and C2) next to the meristem (N=5). **(F’)** Optical section shows *CLE40* expression only in the epidermis of cotyledons. **(G and G’)** MIP **(G)** and optical section **(G’)** of *CLE40* expression (*CLE40:Venus-H2B//Ler*) in a wild type (L.*er*) background at 5WAG shows no signal in the CZ or OC. *CLE40* is confined to the PZ and the centre of older flower primordia, and to sepals (N=8). **(H and H’)** MIP of *CLE40* in a *wus-7* background shows expression through the entire IFM and in the centre of flower primordia. The optical section **(H’)** reveals that *CLE40* is also expressed in the CZ as well as in the OC of the IFM (N= 12). Dashed orange lines indicate the planes of optical sections, dashed blue lines in E and F enclose the meristem, the dashed white line in É marks the CZ. Scale bars: 50µm (A, B, G, H), 20µm (A’, B’, E, E’, F, F’, G’, H’), 1mm (C, D), 500µm (D’), MIP = Maximum intensity projection, PI = propidium iodide, L1 = layer 1 projection, C = cotyledon, LP = leaf primordium

We next determined if *CLE40* repression in the CZ can be alleviated in mutants with reduced WUS activity. Since *wus* loss-of-function mutants fail to maintain an active CZ and shoot meristem, we used the hypomorphic *wus-7* allele (Graf et al., 2010; Ma et al., 2019). *wus-7* mutants are developmentally delayed. Furthermore, *wus-7* mutants generate an IFM, but the FMs give rise to sterile flowers that lack inner organs (Fig3-SupplFig.2). We introgressed the *CLE40* reporter into *wus-7*, and found that at 5WAG, all *wus-7* mutants expressed *CLE40* in both the CZ and the OC of the IFM (Fig. *3*G-H’, Fig3-SupplFig.2). Similar to wild type, *CLE40* is only weakly expressed in the young primordia of *wus-7*. Therefore, we conclude that a *WUS*-dependent pathway downregulates *CLE40* in the centre of the IFM during normal development.

### CLE40 signals through BAM1

Given that CLV1 and BAM1 perform partially redundant functions to perceive CLV3 in shoot and floral meristems, we asked if these receptors also contribute in a CLE40 signalling pathway. We therefore generated the translational reporter lines *CLV1:CLV1-GFP* and *BAM1:BAM1-GFP*, and analysed their expression patterns in detail. We observed dynamic changes of *CLV1* expression during the different stages of flower primordia initiation. *CLV1:CLV1-GFP* is continuously expressed in deeper regions of the IFM comprising the OC, and in the meristem periphery where new FMs are initiating (Fig. *4*A). *CLV1* is expressed strongly in cells of the L1 and L2 of incipient organ primordia (P-1, P0), and only in L2 at P1. P2 and P3 show only very faint expression in the L1, but in stages from P4 to P6, *CLV1* expression expands from the L3 into the L2 and L1 (Fig. *4*, P1-P6).

**Fig. 4:**
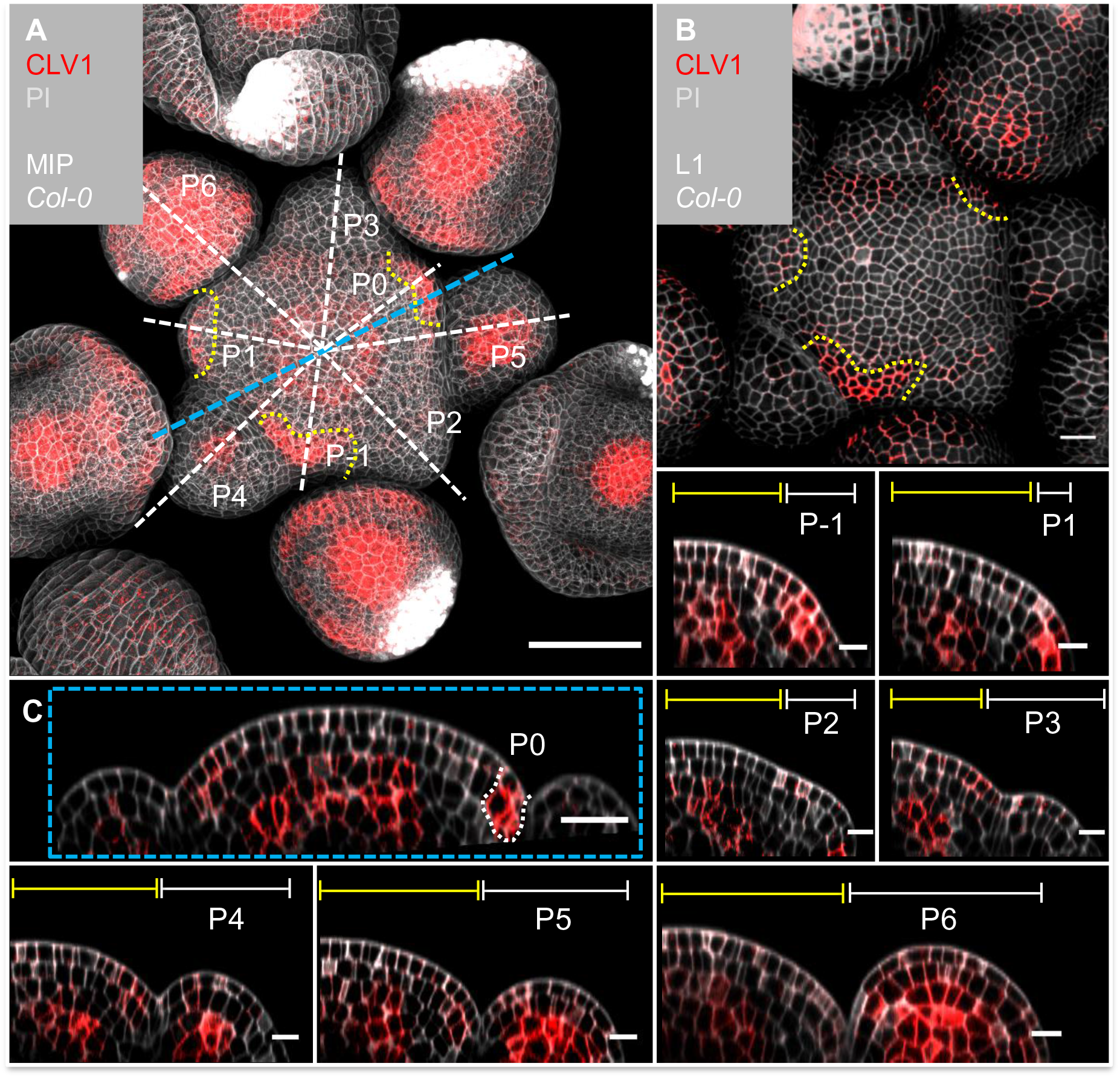
*CLV1* is expressed in the OC and in cells of incipient organ primordia. **(A)** MIP of *CLV1* under its endogenous promoter (*CLV1:CLV1-GFP//Col-0*) at 5 WAG shows *CLV1* expression in the OC of the meristems, IFM and FMs, in incipient organ primordia (P-1 to P1) and in sepals (N=15). **(B)** In the L1 projection *CLV1* expression is detected in cells of incipient organs. **(C)** Optical section through the IFM shows *CLV1* expression in the OC and in P0. **(P-1-P6)** *CLV1* expression is detected in incipient organ primordia in L1 and L2 (-P1, P0), in the L2 of P1, and in the OC of the IFM and FMs from P4 to P6. Dashed white and blue lines indicate the planes of optical sections, yellow dashed line in (A) and (B) mark incipient organ primordia (P-1 to P1), yellow lines (P-1 to P6) indicate the IFM region, white lines mark the primordium. Scale bars: 50µm (A), 20µm (B, C), 10µm (P1 to P6), MIP = maximum intensity projection, PI = propidium iodide, L1 = layer 1, P = primordium

The translational *BAM1:BAM1-GFP* reporter is expressed in the IFM, the FMs and in floral organs (Fig. *5*A). In the IFM, expression is found throughout the L1 layer of the meristem, and, at an elevated level, in L2 and L3 cells of the PZ, but not in the meristem centre around the OC, where *CLV1* expression is detected (Fig. *5*B,C, compare to Fig. *4*C). *BAM1* is less expressed in the deeper regions of primordia from P6 onwards (Fig. *5*C). *BAM1* transcription was reported to be upregulated in the meristem centre in the absence of CLV3 or CLV1 signalling (Nimchuk, 2017). Using our translational BAM1 reporter in the *clv1-20* mutant background, we confirmed that *BAM1* is expressed in the meristem centre, similar to the pattern of *CLV1* in the wild type, and that *BAM1* is upregulated in the L1 of the meristem. Importantly, in a *clv1-20* background BAM1 is absent in the peripheral region of the IFM and the L2 (Fig. *5*D-F).

**Fig. 5:**
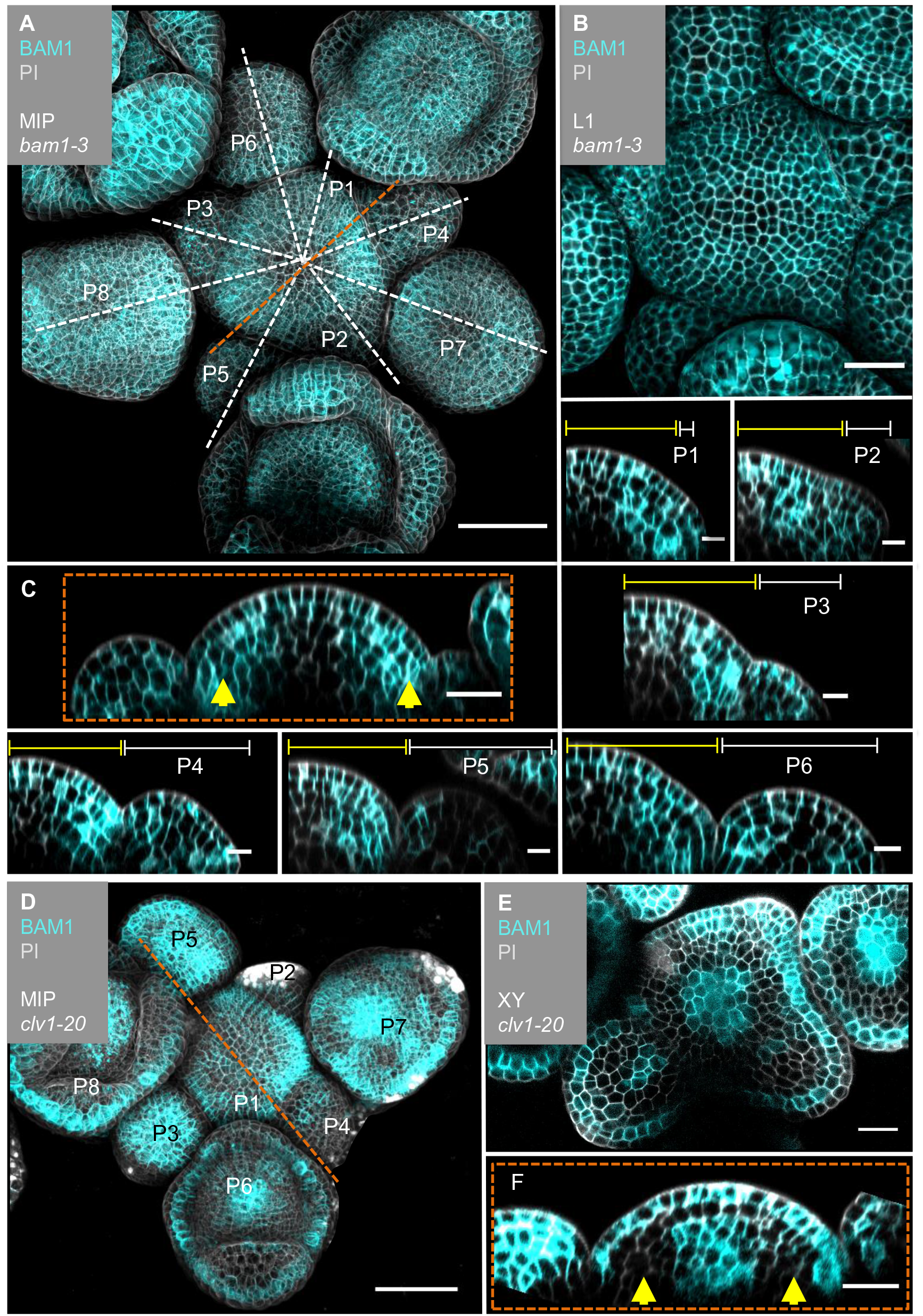
*BAM1* expression is elevated in the flanks of the IFM and not detectable in the OC. **(A)** MIP of *BAM1* under its endogenous promoter (*BAM1:BAM1-GFP//bam1-3*) at 5 WAG. *BAM1* expression is detected nearly throughout the entire inflorescence (IFM, FM, sepals) with weak expression in the CZ of IFM and FMs (N=15). **(B)** The L1 projection of the IFM shows ubiquitous expression of *BAM1*. **(C)** Optical section through the IFM shows elevated *BAM1* expression in the flanks (yellow arrows) and a lack of *BAM1* expression in the OC. **(P1 - P6)** *BAM1* expression is found in all primordia cells. **(D)** MIP of *BAM1* in a *clv1-20* mutant (*BAM1:BAM1-GFP//bam1-3;clv1-20*). *BAM1* expression is detected in most parts of the inflorescence, especially in the centre of the IFM and FMs (N=9). **(E)** Cross section (XY) through of the IFM (from D) shows *BAM1* expression in a *clv1-20* mutant in the CZ (IFM and FMs) and in the L1/L2. **(F)** Optical section through the meristem (from D) shows *BAM1* expression in the OC and in the L1, while no *BAM1* expression is detected in the PZ (yellow arrows). Dashed white and orange lines indicate longitudinal sections; yellow lines (P1 to P6) indicate the IFM region, white lines (P1 to P6) mark the primordium, yellow arrows indicate high (C) or no (F) *BAM1* expression in the PZ. Scale bars: 50µm (A, D), 20µm (B, C, E, F), 10µm (P1 to P6), MIP = maximum intensity projection, PI = propidium iodide, L1 = layer 1, P = primordium

In longitudinal and optical cross sections through the IFM, we found that complementarity of *CLE40* and *CLV3* is reflected in the complementary expression patterns of *BAM1* and *CLV1* (Fig. *6*A-D’). Therefore, we conclude that expression patterns of *CLV1* and *BAM1* are mostly complementary in the meristem itself and during primordia development. When comparing *CLE40* and *BAM1* expression patterns, we found a strong overlap in the peripheral zone of the meristem, during incipient primordia formation, in older primordia, and in L3 cells surrounding the OC (Fig. *6*A’B’, Fig6-SupplFig. 1). Similarly, CLV3 and CLV1 are confined to the CZ and OC, respectively.

**Fig. 6:**
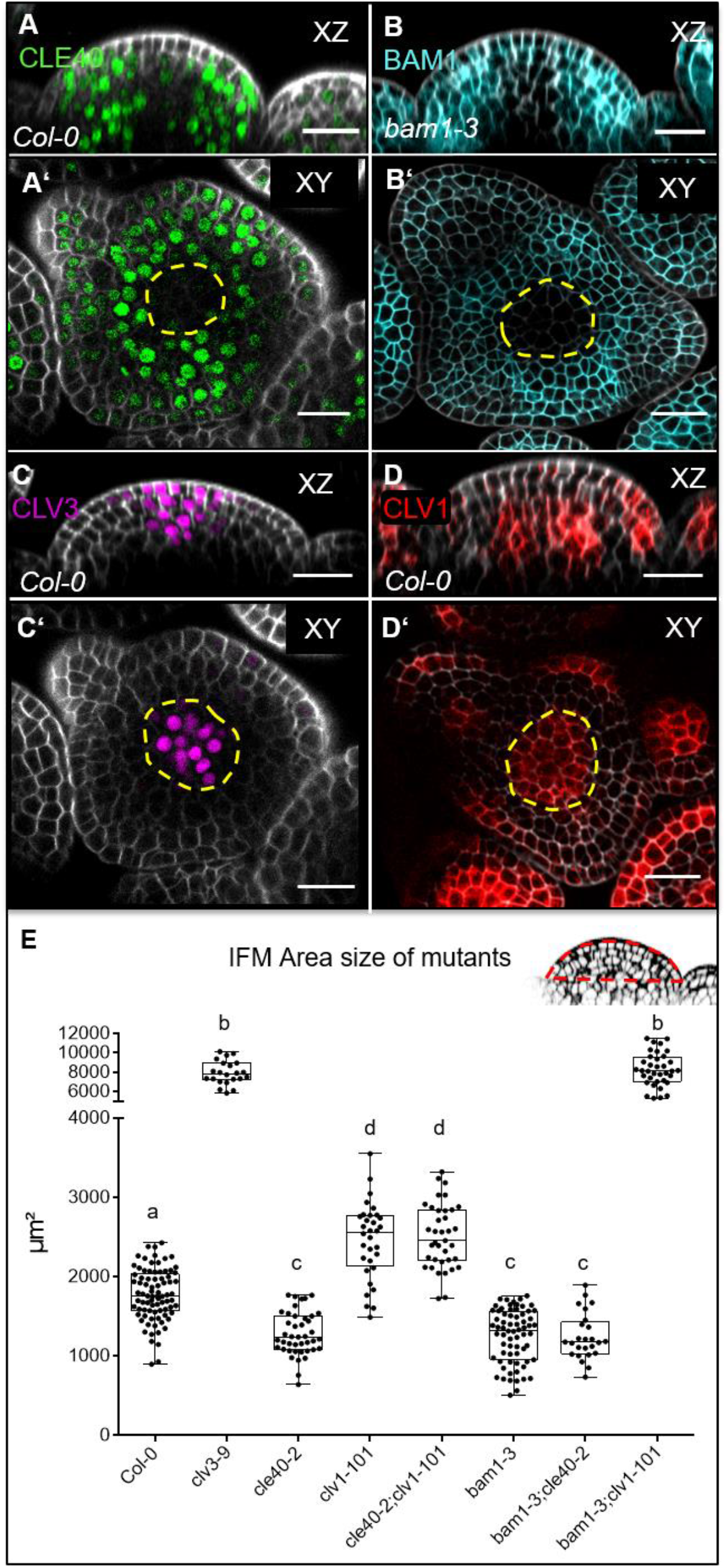
BAM1 and CLV1 are receptors for CLE40 and CLV3, respectively. **(A and A’)** Longitudinal and cross sections of *CLE40* (*CLE40:Venus-H2B*//*Col-0*) through the IFM show *CLE40* expression in the PZ while no *CLE40* expression is detected in the CZ or the OC (dashed yellow line). **(B and B’)** In optical sections of *BAM1* (*BAM1:BAM1-GFP//bam1-3*) through the IFM elevated *BAM1* expression in the PZ and in young primordia can be detected, while low expression is found in the CZ and no expression is observed in the OC (dashed yellow line). **(C and C’)** Optical and cross section of CLV3 through the IFM (*CLV3:NLS-3xmCherry//Col-0*) show *CLV3* expression in the CZ (dashed yellow line). **(D and D’)** The native expression of *CLV1* (*CLV1:CLV1-GFP//Col-0*) in an optical and cross section through the IFM is depicted in the OC (dashed yellow line) and in cells of the L1 and L2 close to emerging primordia. **(E)** Box and whisker plot of the IFM area size of *Col-0* (N=82), various single (*clv3-9* (N=22)*, cle40-2* (N=42)*, clv1-101* (N=32)*, bam1-3* (N=68)) and double mutants (*cle40-2;clv1-101* (N=37)*, cle40-2;bam1-3* (N=25) and *bam1-3;clv1-101* (N=36)) at 6 WAG. Scale bars: 20µm (A – D’), yellow dashed lines indicate the OC (in A’, B’, D’) or the CZ (C’), Statistical groups were assigned after calculating p-values by ANOVA and Turkey’s multiple comparison test (differential grouping from p ≤ 0.01). Red line in the inset meristem in (E) indicates the area that was used for the quantifications in (E).

To analyse if CLE40-dependent signalling requires CLV1 or BAM1, we measured the sizes of IFMs in the respective single and double mutants (Fig. *6*E, Fig6-SupplFig. 2). While *cle40-2* mutant IFMs reached 65% of the wild type size, *clv1-101* plants develop IFMs that were 140% wild type size, whereas *bam1-3;clv1-101* double mutant meristems reached 450% wild type size, similar to those of *clv3-9* mutants. This supports the notion that BAM1 can partially compensate for CLV1 function in the CLV3 signalling pathway when expressed in the meristem centre (Fig. *5*F) (Nimchuk et al., 2015). The relationship between CLV1 and BAM1 is not symmetrical, since CLV1 is expressed in a wildtypic pattern in *bam1-3* mutants (Fig8-SupplFig. 4). Meristem sizes of *bam1-3* mutants reached 70% of the wild type, and double mutants of *cle40-2;bam1-3* did not differ significantly. However, double mutants of *cle40-2;clv1-101* developed like the *clv1-101* single mutant, indicating an epistatic relationship. Importantly, both *clv1-101* and *bam1-3* mutants lack BAM1 function in the meristem periphery (Fig. *5*F), where also *CLE40* is highly expressed, which could explain the observed epistatic relationships of *cle40*-2 with both *clv1-101* and *bam1-3*. Similar genetic relationships for *CLV3, CLE40, CLV1* and *BAM1* were noticed when analysing carpel number as a proxy for FM sizes. We also noted that generation of larger IFMs and FMs in different mutants was negatively correlated with leaf size, which we cannot explain so far (Fig1-SupplFig.1).

We hypothesize that CLE40 signals from the meristem periphery via BAM1 to promote meristem growth. Next, we aimed to determine if the commonalities between *cle40-2* and *bam1-3* mutants extend beyond their effects on meristem size.

### A CLE40 and BAM1 signalling pathway promotes *WUS* expression in the meristem periphery

We next analysed the number of *WUS*-expressing cells in wild type and mutant meristems using a *WUS:NLS-GFP* transcriptional reporter. Compared to wild type, the *WUS* expression domain was laterally strongly expanded in both *clv3-9* and *clv1-101*. Interestingly, *WUS* signal extended also into the L1 layer of *clv1-101*, albeit in a patchy pattern (Fig. *7*A-C’,F). Also noteworthy is that *BAM1* was expressed at a higher level in the L1 layer of *clv1* mutants. *cle40-2* mutants showed a reduction in the number of *WUS* expressing cells down to approx. 50% wild-type levels (Fig. *7*D-D’,F). Importantly, *WUS* remained expressed in the centre of the meristem, but was there found in a narrow domain. In *bam1-3* mutants, the *WUS* domain was similarly reduced as in *cle40-2*, and *WUS* expression focussed in the meristem centre (Fig. *7*E,E’,F). In contrast, both *clv3-9* and *clv1-101* mutants express *WUS* in a laterally expanded domain (Fig. *7*B’,C’).

**Fig. 7:**
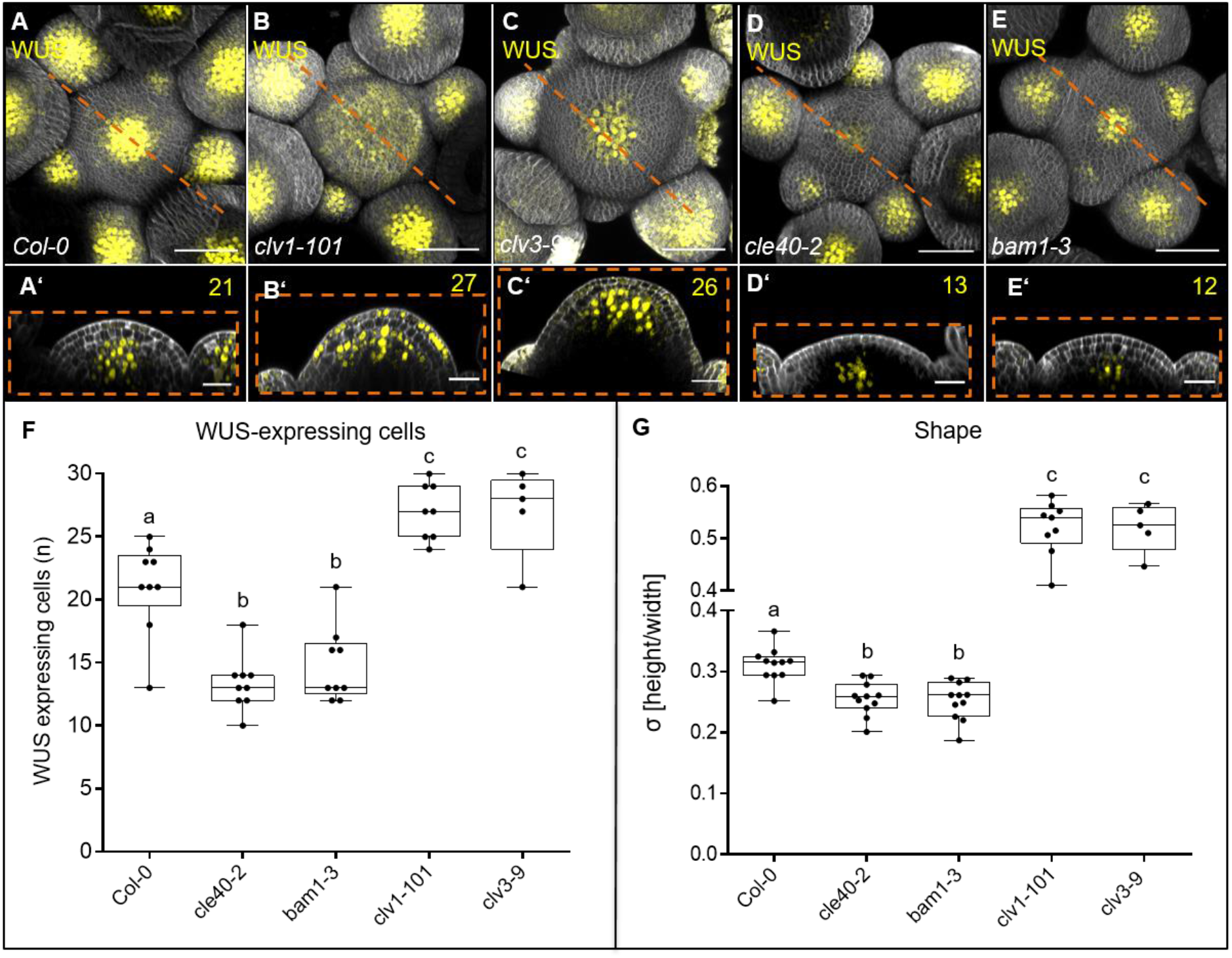
CLE40 and BAM1 promote *WUS* expression. **(A – E’)** MIP and optical section of inflorescences at 5WAG expressing the transcriptional reporter *WUS:NLS-GFP* in a **(A and A’)** *Col-0*, **(B and B’)** *clv1-101*, **(C and C’)** *clv3-9*, **(D and D’)** *cle40-2* and **(E and E’)** *bam1-3* background. In **(A)** wild type plants the WUS domain is smaller compared to the expanded WUS domain in **(B)** *clv1-101* and **(C)** *clv3-9* mutants. The WUS domain of **(D)** *cle40-2* and **(E)** *bam1-3* mutants is decreased compared to wild type plants. Optical sections of **(B’)** *clv1-101* and **(C’)** *clv3-9* mutants expand along the basal-apical axis while the meristem shape of **(D’)** *cle40-2* and **(E’)** *bam1-3* mutants are flatter compared to **(A’)** wild type plants,. **(F)** Box and whisker plot shows the number of *WUS*-expressing cells in the OC of IFMs of *Col-0* (N=9), *cle40-2* (N=9), *bam1-3* (N=9), *clv1-101* (N=8), and *clv3-9* (N=5). **(G)** After 5 WAG *bam1-3* (N=11) and *cle40-2* (N=11) mutants have flatter meristems than wild type plants (decreased σ value compared to *Col-0* (N=11)), while *clv1-101* (N=9) and *clv3-9* (N=6) mutants increase in their IFM height showing a higher σ value. Scale bars: 50µm (A – E), 20µm (A’-J), Statistical groups and stars were assigned after calculating p-values by ANOVA and Turkey’s multiple comparison test (differential grouping from p ≤ 0.01). WAG = weeks after germination, yellow numbers = *WUS* expressing cells in the CZ, σ value = height/width of IFMs

To integrate our finding that *CLE40* expression is repressed by WUS activity with the observation that *WUS*, in turn, is promoted by *CLE40* signalling, we hypothesize that the *CLE40-BAM1-WUS* interaction establishes a new negative feedback loop. The *CLE40-BAM1-WUS* negative feedback loop acts in the meristem periphery, while the *CLV3-CLV1-WUS* negative feedback loop acts in the meristem centre along the apical-basal axis. Both pathways act in parallel during development to regulate the size of the *WUS* expression domain in the meristem, possibly by perceiving input signals from two different regions, the CZ and the PZ, of the meristem.

We then asked how the two signalling pathways, converge on the regulation of *WUS* expression, control meristem growth and development. So far, we showed that both *CLV3-CLV1* and *CLE40-BAM1* signalling control meristem size, but in an antagonistic manner. However, we noticed that the different mutations in peptides and receptors affected distinct aspects of meristem shape. We therefore analysed meristem shape by measuring meristem height (the apical-basal axis) at its centre, and meristem diameter (the radial axis) at the base in longitudinal sections. The ratio of height to width then gives a shape parameter “σ” (from the greek word σχήμα = shape). In young inflorescence meristems at 4-5 WAG, when inflorescence stems were approximately 5-8 cm long, meristems of *cle40-2* and *bam1-3* mutants were slightly reduced in width, and strongly reduced in height, resulting in reduced σ in comparison to *Col-0* (Fig. *7*G, Fig7-SupplFig. 1A). Meristems of *clv1-101* and *clv3-9* mutants were similar in width to wild type, but strongly increased in height, giving high σ values (Fig. *7*, Fig7-SupplFig. 1A-C). This indicates that *CLV3-CLV1* signalling mostly restricts meristem growth along the apical-basal axis, while *CLE40-BAM1* signalling promotes meristem growth along both axes.

Our data expand the current model of shoot meristem homeostasis by taking into account that stem cells are lost from the OC during organ initiation in the PZ (Fig. *8*). CLV3 signals from the CZ via CLV1 in the meristem centre to confine *WUS* expression to the OC. The diffusion of WUS protein along the apical-basal axis towards the meristem tip establishes the CZ and activates *CLV3* expression as a feedback signal. During plant growth, rapid cell division activity and organ initiation requires the replenishment of PZ cells from the CZ, which can be mediated by increased *WUS* activity. We now propose that the PZ generates CLE40 as a short range or autocrine signal that acts through BAM1 in the meristem periphery. Since *BAM1* and *WUS* expression do not overlap, we postulate the generation of a diffusible factor that relies on *CLE40-BAM1*, and acts from the PZ to promote *WUS* expression. WUS, in turn, represses *CLE40* expression from the OC, thus establishing a second negative feedback regulation. Together, the two intertwined pathways serve to adjust WUS activity in the OC and incorporate information on the actual size of the stem cell domain, via *CLV3-CLV1*, and the growth requirements from the PZ via *CLE40-BAM1*.

**Fig. 8:**
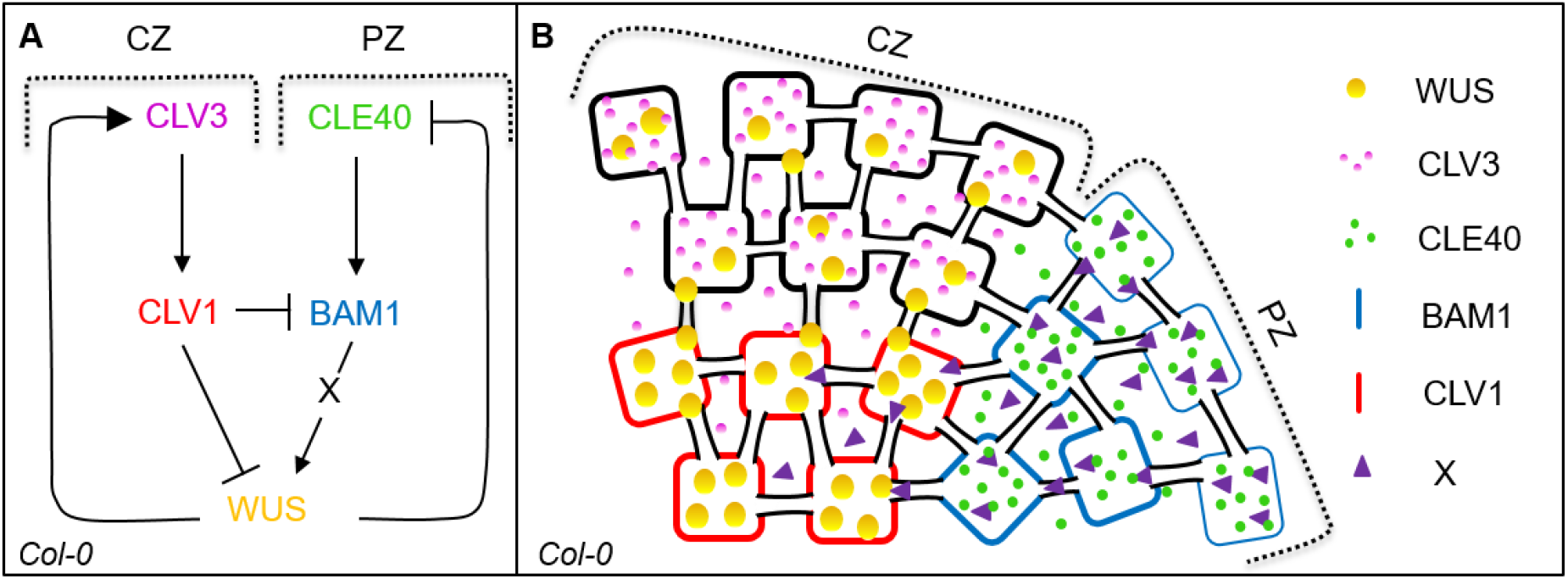
Schematic model of two intertwined signalling pathways in the shoot meristem. **(A and B)** Schematic representation of two negative feedback loops in the IFM of *Arabidopsis thaliana*. CLV3 in the CZ binds to the LRR receptor CLV1 to activate a downstream signalling cascade which leads to the repression of the transcription factor WUS. In a negative feedback loop WUS protein moves to the stem cells to activate *CLV3* gene expression. In the PZ of the IFM a second negative feedback loop controls meristem growth by CLE40 and its receptor BAM1. CLE40 binds to BAM1 in an autocrine manner, leading to the activation of a downstream signal “X” which promotes WUS activity. WUS protein in turn represses the expression of the *CLE40* gene. CZ = central zone, PZ = peripheral zone, arrows indicate a promoting effect and the blocked line indicates a repressing signal

## Discussion

Shoot meristems are the centres of growth and organ production throughout the life of a plant. Meristems fulfil two main tasks, which are the maintenance of a non-differentiating stem cell pool, and the assignment of stem cell daughters to lateral organ primordia and differentiation pathways (Hall & Watt, 1989). Shoot meristem homeostasis requires extensive communication between the CZ, the OC and the PZ. The discovery of CLV3 as a signalling peptide, which is secreted exclusively from stem cells in the CZ, and its interaction with WUS in a negative feedback loop was fundamental for our understanding of such communication pathways (Fletcher, 2020). Here, we analysed the function of *CLE40* in shoot development of Arabidopsis, and found that *WUS* expression in the OC is under positive control from the PZ due to the activity of a CLE40-BAM1 signalling pathway. IFM size is reduced in *cle40* mutants, indicating that *CLE40* signalling promotes meristem size. Importantly, *CLE40* is expressed in the PZ, in late stage FMs and in differentiating organs. A common denominator for the complex and dynamic expression pattern is that *CLE40* expression is confined to meristematic tissues, but not in organ founder sites or in regions with high WUS activity, such as the OC and the CZ. Both misexpression of *WUS* in the *CLV3* domain (Fig. *3*F), studies of *clv3* mutants with expanded stem cell domains (Fig. *3*B, Fig3-SupplFig.1) and analysis of *wus* mutants (Fig. *3*, Fig3-SupplFig.2) underpinned the notion that *CLE40* expression, in contrast to *CLV3*, is negatively controlled in a *WUS*-dependent manner. Furthermore, we found that the number of *WUS* expressing cells in *cle40* mutant IFMs is strongly reduced, indicating that *CLE40* exerts its positive effects on IFM size by expanding the *WUS* expression domain.

So far, the antagonistic effects of Arabidopsis *CLV3* and *CLE40* on meristem size can only be compared to the antagonistic functions of *MpCLE1* and *MpCLE2* on the gametophytic meristems of *M. polymorpha*, which signal through two distinct receptors, MpTDR and MpCLV1, respectively (Hata & Kyozuka, 2021). By the complementation of *clv3* mutants through expression of *CLE40* from the *CLV3* promoter it was shown previously that *CLE40* and *CLV3* are able to activate the same downstream receptors (Hobe et al., 2003). Our detailed analysis of candidate receptor expression patterns showed that *CLV3* and *CLV1* are expressed in partially overlapping domains in the meristem centre, while *CLE40* and *BAM1* are confined to the meristem periphery. Like *cle40* mutants, *bam1* mutant IFMs are smaller and maintain a smaller *WUS* expression domain, supporting the notion that *CLE40* and *BAM1* comprise a signalling unit that increases meristem size by promoting *WUS* expression. The antagonistic functions of the *CLV3-CLV1* and *CLE40-BAM1* pathways in the regulation of *WUS* are reflected in their complementary expression patterns. There is cross-regulation between these two signalling pathways at two levels: (1) *WUS* has been previously shown to promote *CLV3* levels in the CZ, and we here show that *WUS* represses (directly or indirectly) *CLE40* expression in the OC and in the CZ (Fig. *3*B, Fig3-SupplFig.1); (2) *CLV1* represses *BAM1* expression in the OC, and thereby restricts *BAM1* to the meristem periphery (Fig. *5*, Fig. *6*). In *clv1* mutants, *BAM1* shifts from the meristem periphery to the OC, and the *WUS* domain laterally expands in the meristem centre (Fig. *5*F, Fig. *7*B’). Furthermore, *BAM1* expression increases also in the L1, which could cause the observed irregular expression of *WUS* in the outermost cell layer of *clv1* mutants. The role of BAM1 in the OC is not entirely clear: despite the high sequence similarity between CLV1 and BAM1, the expression of *BAM1* in the OC is not sufficient to compensate for the loss of CLV1 (Fig.5D-F, Nimchuk et al., 2015). In the OC, BAM1 appears to restrict *WUS* expression to some extent, since *clv1;bam1* double mutants reveal a drastically expanded IFM (DeYoung & Clark, 2008). However, it is possible that BAM1 in the absence of CLV1 executes a dual function: to repress *WUS* in response to CLV3 in the OC as a substitute for CLV1, and simultaneously to promote *WUS* expression in the L1 in response to CLE40.

The expression domains of *CLE40* and its receptor *BAM1* largely coincide, suggesting that CLE40 acts as an autocrine signal. Similarly, protophloem sieve element differentiation in roots is inhibited by CLE45, which acts as an autocrine signal via BAM3 (Kang & Hardtke, 2016). Since *WUS* is not expressed in the same cells as *BAM1*, we have to postulate a non-cell autonomous signal X that is generated in the peripheral zone due to *CLE40-BAM1* signalling, and diffuses towards the meristem centre to promote *WUS* expression (Hohm et al., 2010). As a result, *CLE40-BAM1* signalling from the PZ will provide the necessary feedback signal that stimulates stem cell activity and thereby serves to replenish cells in the meristem for the initiation of new organs. The *CLV3-CLV1* signalling pathway then adopts the role of a necessary feedback signal that avoids an excessive stem cell production.

The two intertwined, antagonistically acting signalling pathways that we described here allow us to better understand the regulation of shoot meristem growth, development and shape. The previous model, which focussed mainly on the interaction of the CZ and the OC via the *CLV3-CLV1-WUS* negative feedback regulation, lacked any direct regulatory contribution from the PZ. *EPFL* peptides were shown to be expressed in the periphery and to restrict both *CLV3* and *WUS* expression via ER (Zhang et al., 2021). However, *EPFL* peptide expression is not reported to be feedback regulated from the OC or CZ, and the main function of the *EPFL-ER* pathway is therefore to restrict overall meristem size (Zhang et al., 2021). The second negative feedback loop controlled by CLE40, which we uncovered here, enables the meristem to fine-tune stem cell activities in response to fluctuating requirements for new cells during organ initiation. Due to the combined activities of CLV3 and CLE40, the OC (with WUS as a key player) can now record and compute information from both, the CZ and PZ. Weaker *CLV3* signalling, indicating a reduction in the size of the CZ, induces preferential growth of the meristem along the apical-basal axis (increasing σ), while weaker CLE40 signals, reporting a smaller PZ, would decrease σ and flatten meristem shape. It will be intriguing to investigate if different levels of CLV3 and CLE40 also contribute to the shape changes that are observed during early vegetative development, or upon floral transition in Arabidopsis.

Many shoot-expressed CLE peptides are encoded in the genomes of maize, rice and barley, which could act analogously to CLV3 and CLE40 of Arabidopsis. It is tempting to speculate that in grasses, a CLE40-like, stem cell promoting signalling pathway is more active than a CLV3-like, stem cell restricting pathway. This could contribute to the typical shape of cereal SAMs, which are, compared to the dome-shaped SAM of dicotyledonous plants, extended along the apical-basal axis.

## Material and Methods

All chemicals used for the experiments are listed in Tab. *1*.

**Tab. 1:** Chemicals and substances used in this study.

| Name | Producer | Product no. | CAS no. |
| --- | --- | --- | --- |
| BASTA® non-selective herbicide | Bayer CropScience | 84442615 | N/A |
| bacto™ agar | gibco | 214010 | 9105960 |
| bacto™ yeast extract | gibco | 212750 | 9070604 |
| Carbenicillin disodium salt | Carl Roth | 6344.2 | 4800-94-6 |
| DAPI (4,6-diamidino-2-phenylindole) | N/A | N/A | 28718-90-3 |
| DL-phosphinothricin (PPT) | Duchefa Biochemie bv | P0159 | 77182-82-2 |
| D-Alanin | Sigma-Aldrich (Merck) | A7377 | 338-69-2 |
| D(+)-Saccharose | Carl Roth | 4661.1 | N/A |
| Gentamicin sulfate | Sigma-Aldrich (Merck) | G1264 | 1405-41-0 |
| Hygromycin B | Duchefa Biochemie bv | H0192 | 31282-04-9 |
| Hypochloride acid (~37%) | Thermo Fischer Scientific | H/120/PB15 | 1884567 |
| Kanamycin monosulfate | Duchefa Biochemie bv | K0126.0005 | 25389-94-0 |
| Magnesium chloride x 6H<sub>2</sub>O | Grüssig GmbH | 12087 | 205 |
| MES hydrate | Sigma-Aldrich (Merck) | 10240885 | 1266615-59-1 |
| Murashige & Skoog<br>(+ Gamborg B5 vitamins) | Duchefa Biochemie bv | M0231.0050 | N/A |
| Phusion High-Fidelity PCR<br>polymerase | Thermo Fischer Scientific | F530S | N/A |
| Plant agar | Duchefa Biochemie bv | P1001.1000 | 9002-18-0 |
| Potassium hydroxide | Sigma-Aldrich (Merck) | 9643807 | N/A |
| Propidium iodide | Thermo Fischer Scientific | P1304MP | 25535-16-4 |
| Rifampicin | TCI | R0079 | 13292-46-1 |
| Spectinomycin HCl pentahydrate | Duchefa Biochemie bv | S0188 | 22189-32-8 |
| Sodium dodecyl sulfate (SDS) | Sigma-Aldrich (Merck) | L3771 | 151-21-3 |
| Sodium chloride | Carl Roth | 3957.1 | N/A |
| Sodium hypochloride (13%) | Zentrale Chemikalienlager (ZCL) | 2N370 | N/A |
| Tetracycline | Sigma-Aldrich (Merck) | 87128 | 60-54-8 |
| Tryptone | gibco | N/A | 57091 |

### Plant material and growth conditions

All wild type *Arabidopsis thaliana* (L.) Heynh. plants used in this study are ecotype Columbia-0 (*Col-0*), except for *wus-7* mutants which are in Landsberg *erecta* (L.*er*.) background. Details about *Arabidopsis thaliana* plants carrying mutations in the following alleles: *bam1-3, cle40-2, cle40-cr1, cle40-cr2, cle40-cr3, clv1-101, clv3-9* and *wus-7* are described in Tab. *2*. All mutants are in *Col-0* background and are assumed to be null-mutants, except for *wus-7* mutants. *cle40* mutants (*cle40-2, cle40-cr1, cle40-cr2, cle40-cr3*) have either a stop codon, a T-DNA insertion or deletion in or before the crucial CLE box domain (Fig1-SupplFig.2B’). *clv3-9* mutants were generated in 2003 by the lab of R. Simon. *clv3-9* mutants were created by EMS resulting in a W62STOP mutation before the critical CLE domain region. *bam1-3* and *clv1-101* mutants have been described as null mutants before (DeYoung et al., 2006; Kinoshita et al., 2010), while *clv1-20* is a weak allele which contains a insertion within the 5’-UTR of CLV1 and results in a reduced mRNA level (Durbak & Tax, 2011). *wus-7* is a weak allele and mutants were described in previous publications (Graf et al., 2010). Double mutants were obtained by crossing the single mutant plants until both mutations were proven to be homozygous for both alleles. Genotyping of the plants was performed either by PCR or dCAPS method with the primers and restrictions enzymes listed in Tab. *3*. Before sowing, seeds were either sterilized for 10min in an ethanol solution (80% v/v ethanol, 1,3% w/v sodium hypochloride, 0,02% w/v SDS) or for 1h in a desiccator in a chloric gas atmosphere (50mL of 13% w/v sodium hypochlorite with 1mL 37% HCL). Afterwards, seeds were stratified for 48h at 4°C in darkness. Seeds on soil were then cultivated in phytochambers under long day (LD) conditions (16h light/ 8h dark) at 21°C. For selection of seeds or imaging of vegetative meristems seeds were sowed on ½ Murashige & Skoog (MS) media (1% w/v sucrose, 0.22% w/v MS salts + B5 vitamins, 0.05% w/v MES, 12g/L plant agar, adjusted to pH 5.7 with KOH) in squared petri dishes. Seeds in petri dishes were kept in phytocabinets under continuous light conditions at 21°C and 60% humidity.

**Tab. 2:** Mutants analysed in this study.

| **Allele** | **Gene** | **Mutation** | **Reference** | **Background** |
| --- | --- | --- | --- | --- |
| *bam1-3* | AT5G65700 | T-DNA | Alonso et al., 2003;<br>SALK_015302 | *Col-0* |
| *cle40-2* | AT5G12990 | Transposon<br>mutation | Stahl et al., 2009 | *Col-0* |
| *cle40-cr1*<br>*cle40-cr2*<br>*cle40-cr3* | AT5G12990 | CRISPR | Yamaguchi et al., 2017 | *Col-0* |
| *clv3-9* | AT2G27250 | EMS | Rüdiger Simon, 2003 | *Col-0* |
| *clv1-20* | AT1G75820 | T-DNA | SALK_008670 | *Col-0* |
| *clv1-101* | AT1G75820 | T-DNA | Kinoshita et al., 2010;<br>CS858348 | *Col-0* |
| *wus-7* | AT2G17950 | EMS | Graf et al., 2010 | *L.er.* |

**Tab. 3:**
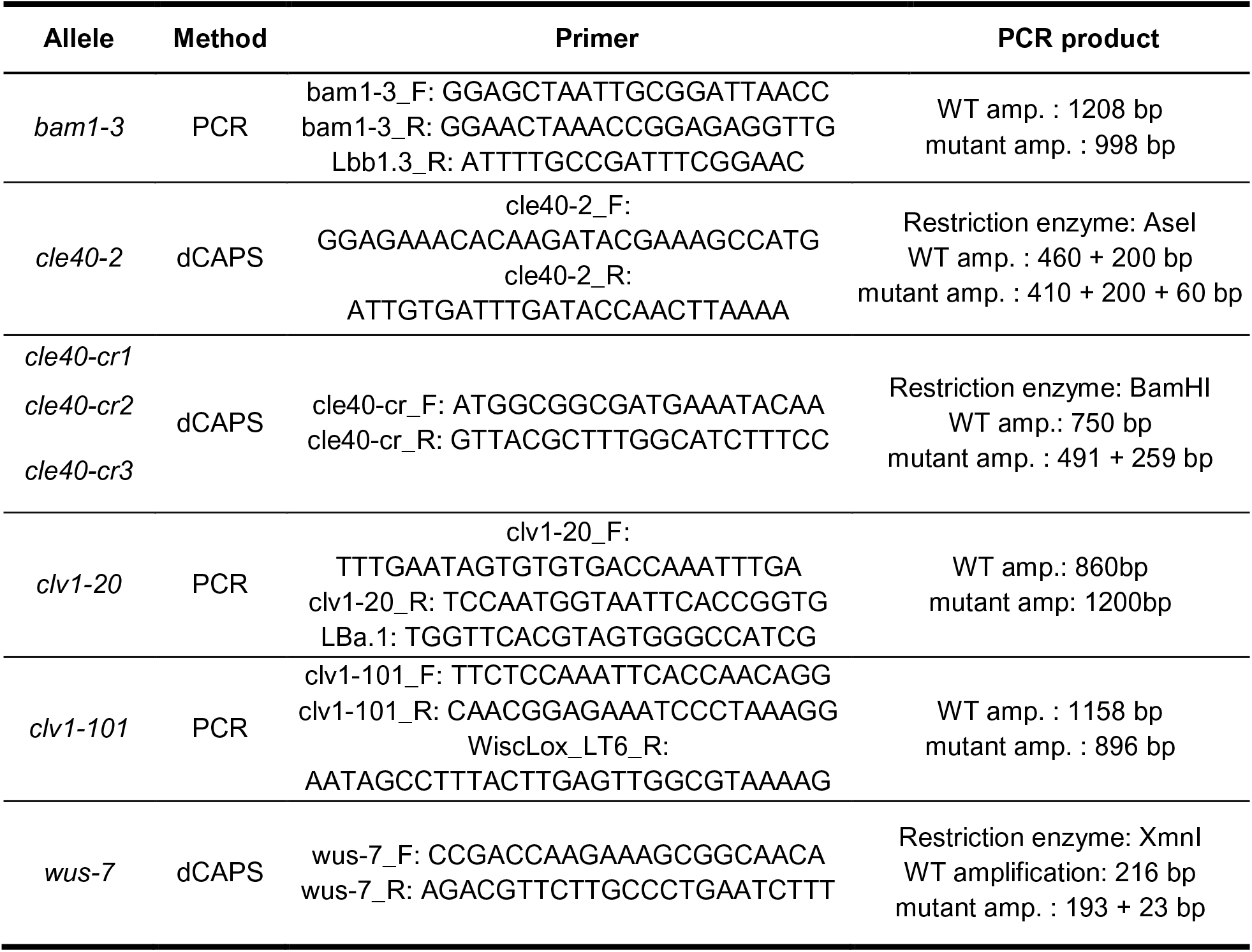
Primers and methods used for genotyping.

### Cloning of reporter lines

CLV1 (*CLV1:CLV1-GFP*), BAM1 (*BAM1:BAM1-GFP*) and CLV3 (*CLV3:NLS-3xmCherry*) reporter lines were cloned using the GreenGate method (Lampropoulos et al., 2013). Entry and destination plasmids are listed in Tab. *4* and Tab. *5*. Promoter and coding sequences were PCR amplified from genomic *Col-0* DNA which was extracted from rosette leaves of *Col-0* plants. Primers used for amplification of promoters and coding sequences can be found in Tab. *6* with the specific overhangs used for the GreenGate cloning system. Coding sequences were amplified without the stop codon to allow transcription of fluorophores at the C-terminus. BsaI restriction sites were removed by site-directed mutagenesis using the “QuickChange II Kit” following the manufacturer’s instructions (Agilent Technologies). Plasmid DNA amplification was performed by heat-shock transformation into *Escherichia coli* DH5α cells (10min on ice, 1min at 42°C, 1min on ice, 1h shaking at 37°C), which were subsequently plated on selective LB medium (1% w/v tryptone, 0.5% w/v yeast extract, 0.5% w/v NaCl) and cultivated overnight at 37°C. All entry and destination plasmids were validated by restriction digest and Sanger sequencing.

**Tab. 4:** Entry vectors used for cloning.

| Name | Description | Bacterial resistance | Backbone | Reference/ Origin |
| --- | --- | --- | --- | --- |
| proBAM1 (pGD288) | BAM1 promoter<br>3522bp upstream from transcription start | Ampicillin | pGGA000 | Grégoire Denay |
| proCLV3 | CLV3 promoter<br>1480bp upstream from transcription start | Ampicillin | pGGA000 | Jenia Schlegel |
| proCLV1 | CLV1 promoter<br>5759bp upstream from transcription start | Ampicillin | pGGA000 | Patrick Schultz |
| omega-element (pGGB002) | Omega- element | Ampicillin | pGGB000 | Lampropoulos et al., 2013 |
| SV40 NLS (pGGB005) | SV40 NLS (SIMIAN VIRUS 40 NUCLEAR LOCALIZATION SIGNAL) | Ampicillin | pGGB000 | Lampropoulos et al., 2013 |
| BAM1_CDS (pGD351) | BAM1 coding region<br>genomic region of BAM1 START to one codon before STOP, including introns, internal Bsal sites removed | Ampicillin | pGGC000 | Grégoire Denay |
| CLV1_CDS | CLV1 coding region<br>2946 bp coding region amplified from genomic Col-0 DNA without STOP codon and internal Bsal site removed | Ampicillin | pGGC000 | Jenia Schlegel |
| 3x-mCherry (pGGC026) | 3x mCherry | Ampicillin | pGGC000 | Lampropoulos et al., 2013 |
| linker-GFP (pGD165) | linker(10aa)-eGFP | Ampicillin | pGGD000 | Grégoire Denay |
| d-dummy (pGGD002) | d-dummy | Ampicillin | pGGD000 | Lampropoulos et al., 2013 |
| tCLV3 | CLV3 terminator<br>1257 bp downstream of transcription stop | Ampicillin | pGGE000 | Jenia Schlegel |
| tUBQ10 (pGGE009) | UBQ10 terminator | Ampicillin | pGGE000 | Lampropoulos et al., 2013 |
| BastaR (pGGF008) | pNOS:BastaR (chi sequence removed):tNOS | Ampicillin | pGGF000 | Lampropoulos et al., 2013 |
| D-AlaR (pGGF003) | pMAS:D-AlaR:tMAS | Ampicillin | pGGF000 | Lampropoulos et al., 2013 |

**Tab. 5:** Destination vectors used to generate transgenic *A.thaliana* reporter lines.

| Name | Backbone | Promoter | N-tag | CDS | C-tag | Terminator | Resistance |
| --- | --- | --- | --- | --- | --- | --- | --- |
| *BAM1:BAM1-GFP* | pGGZ001 | proBAM1 | Ω- element (pGGB002) | BAM1-CDS | linker-GFP (pGD165) | tUBQ10 (pGGE009) | D-Alanin (pGGF003) |
| *CLV1:CLV1-GFP* | pGGZ001 | proCLV1 | Ω- element (pGGB002) | CLV1-CDS | linker-GFP (pGD165) | tUBQ10 (pGGE009) | BastaR (pGGF008) |
| *CLV3:NLS-3xmCherry* | pGGZ001 | proCLV3 | SV40 NLS | 3x-mCherry (pGGC026) | d-dummy (pGGD002) | tCLV3 | BastaR (pGGF008) |

**Tab. 6:**
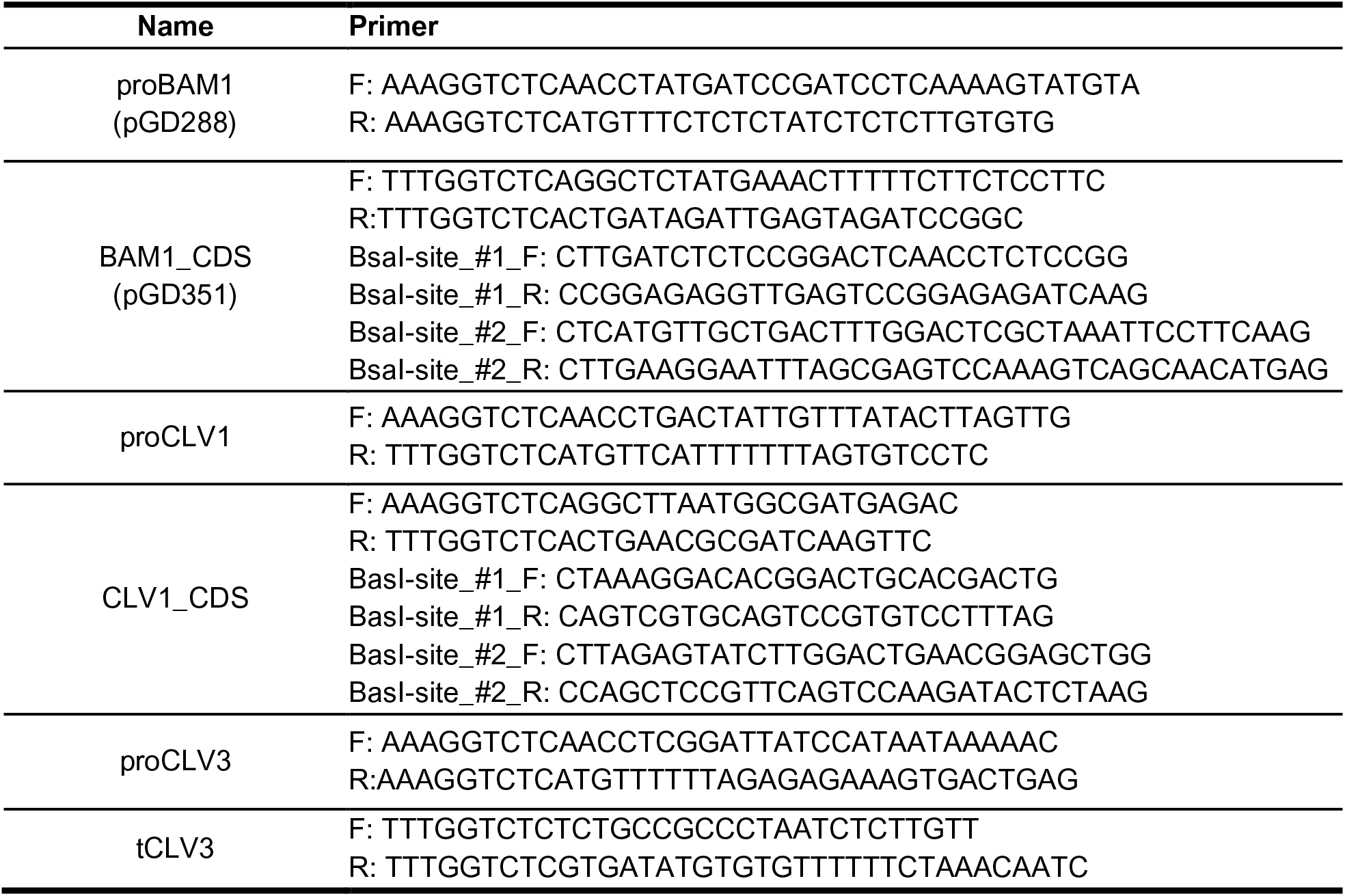
Primers used for cloning the entry vectors.

### Generation of stable *A. thaliana* lines

Generation of stable *Arabidopsis thaliana* lines was done by using the floral dip method (Clough & Bent, 1998).

Translational *CLV1* (*CLV1:CLV1-GFP*) and transcriptional *CLV3* (*CLV3:NLS-3xmCherry*) reporter carry the BASTA plant resistance cassette. T1 seeds were sown on soil and sprayed with Basta ® (120mg/mL) after 5 and 10 DAG. Seeds of ∼10 independent Basta-resistant lines were harvested. The translational *BAM1* (*BAM1:BAM1-GFP*) reporter line carries a D-Alanin resistance cassette and T1 seeds were sown on ½ MS media containing 3-4mM D-Alanin. T2 seeds were then selected on ½ MS media supplied with either 3-4mM D-Alanin or 10µg/mL of DL-phosphinothricin (PPT) as a BASTA alternative. Only plants from lines showing about ∼75% viability were kept and cultivated under normal plant conditions (21°C, LD). Last, T3 seeds were plated on ½ MS media supplied with 3-4mM D-Alanin or PPT again and plant lines showing 100% viability were kept as homozygous lines. The *CLV3:NLS-3xmCherry* and *CLV1:CLV1-GFP* constructs were transformed into *Col-0* wild type plants and after a stable T3 line was achieved, plants carrying the *CLV1:CLV1-GFP* construct were crossed into *bam1-3, cle40-2, clv3-9* and *clv1-101* mutants until a homozygous mutant background was reached. *BAM1:BAM1-GFP* lines were floral dipped into *bam1-3* mutants and subsequently crossed into the *clv1-20* mutant background which rescued the extremely fasciated meristem phenotype of *bam1-3;clv1-20* double mutants (Fig. *5*D-F). *BAM1:BAM1-GFP//bam1-3* plants were also crossed into *cle40-2* and *clv3-9* mutants until a homozygous mutant background was achieved. The *CLE40:Venus-H2B* reporter line was created and described in Wink, 2013 and the *WUS:NLS-GFP;CLV3:NLS-mCherry* reporter line was a gift from the Lohmann lab (Wink, 2013). *CLE40:Venus-H2B* reporter line was crossed into homozygous *clv3-9* and heterozygous *wus-7* mutants. Homozygous *clv3-9* mutants were detected by its obvious phenotype and were brought into a stable F3 generation. Homozygous *wus-7* mutants were genotyped. Seeds were kept in the F2 generation, since homozygous *wus-7* plants do not develop seeds. The *CLE40:Venus-H2B* reporter line was also crossed with the *CLV3:NLS-3xmCherry* reporter line and was brought into a stable F3 generation. To generate the *CLE40:Venus-H2B//CLV3:WUS* line, plants carrying the *CLE40:Venus-H2B* line were floral dipped with the *CLV3:WUS* construct. T1 seeds were sown on 10µg/mL of DL-phosphinothricin (PPT) and the viable seedlings were imaged. *WUS:NLS-GFP/CLV3:NLS-mCherry//Col-0* reporter line was crossed into *clv3-9, cle40-2, clv1-101* and *bam1-3* mutants until a stable homozygous F3 generation was reached respectively. Detailed information of all used *A. thaliana* lines can be found in Tab. *7*.

**Tab. 7:** Arabidopsis lines that were analysed in this study.

| Name/Construct | Background | Plant resistance | Generation | Reference |
| --- | --- | --- | --- | --- |
| *BAM1:BAM1-GFP* | *bam1-3* | D-Ala | T4 | this study |
| *BAM1:BAM1-GFP* | *bam1-3;clv1-20* | D-Ala | F3 | this study |
| *BAM1:BAM1-GFP* | *bam1-3;clv3-9* | D-Ala | F3 | this study |
| *BAM1:BAM1-GFP* | *bam1-3;cle40-2* | D-Ala | F3 | this study |
| *CLE40:Venus-H2B* | *Col-0* | Hygromycin | T5 | Rene Wink |
| *CLE40:Venus-H2B* | *clv3-9* | Hygromycin | F3 | this study |
| *CLE40:Venus-H2B* | *wus-7* | Hygromycin | F2 | this study |
| *CLE40:Venus-H2B* | *CLV3:WUS//Col-0* | Hygromycin/<br>Basta | T1* | this study |
| *CLV1:CLV1-GFP* | *Col-0* | Basta | T4 | this study |
| *CLV1:CLV1-GFP* | *bam1-3* | Basta | F3 | this study |
| *CLV1:CLV1-GFP* | *clv3-9* | Basta | F3 | this study |
| *CLV1:CLV1-GFP* | *cle40-2* | Basta | F3 | this study |
| *CLV3:NLS-3xmCherry* | *CLE40:Venus-H2B//Col-0* | Basta/<br>Hygromycin | F3 | this study |
| *CLV3:NLS-mCherry*<br>*WUS:NLS-GFP* | *Col-0* | Kanamycin | N/A | Anne Pfeiffer |
| *CLV3:NLS-mCherry*<br>*WUS:NLS-GFP* | *cle40-2* | Kanamycin | F3 | this study |
| *CLV3:NLS-mCherry*<br>*WUS:NLS-GFP* | *bam1-3* | Kanamycin | F3 | this study |
| *CLV3:NLS-mCherry*<br>*WUS:NLS-GFP* | *clv1-101* | Kanamycin | F3 | this study |
| *CLV3:NLS-mCherry*<br>*WUS:NLS-GFP* | *clv3-9* | Kanamycin | F3 | this study |

\* Plants do not overcome seedling stage

### Confocal imaging of IFMs

To image IFMs *in vivo*, plants were grown under LD (16h light/ 8h dark) conditions and inflorescences were cut off at 5 or 6 WAG. Inflorescences were stuck on double sided adhesive tape on an objective slide and dissected until only the meristem and primordia from P0 to maximum P10 were visible. Next, inflorescences were stained with either propidium iodide (PI 5mM) or 4′,6-Diamidin-2-phenylindol (DAPI 1µg/mL) for 2 to 5min. Inflorescences were then washed three times with water and subsequently covered with water and a cover slide and placed under the microscope. Imaging was performed with a Zeiss LSM780 or LSM880 using a W Plan-Apochromat 40x/1.2 objective. Laser excitation, emission detection range and detector information for fluorophores and staining can be found in Tab. *8*. All IFMs were imaged from the top taking XY images along the Z axis, resulting in a Z-stack through the inflorescence. The vegetative meristems were imaged as described for IFMs. Live imaging of the reporter lines in *A. thaliana* plants was performed by dissecting primary inflorescences (except for *clv3-9* mutants) at 5 WAG under LD conditions. For imaging of the reporter lines in the mutant backgrounds of *clv3-9* secondary IFMs were dissected, since the primary meristems are highly fasciated. Vegetative meristems were cultivated in continuous light conditions at 21°C on ½ MS media plates and were imaged at 10 DAG. For each reporter line at least 3 independent experiments were performed and at least 5 IFMs were imaged.

**Tab. 8:** Microscopy settings used for imaging.

| Fluorophore/<br>Staining | Excitation | Emission | MBS | Detector | Light source |
| --- | --- | --- | --- | --- | --- |
| DAPI | 405 nm | 410 - 490 nm | 405 | PMT* | Diode |
| GFP | 488 nm | 500 - 545 nm | 488/561 | GaAsP | Argon laser |
| Venus | 514 nm | 518 - 540 nm | 458/514 | GaAsP | Argon laser |
| mCherry | 561 nm | 570 - 640 nm | 458/561 | PMT* | DPSS laser** |
| PI | 561 nm | 595 - 650 nm | 488/561 | PMT* | DPSS laser** |

\* Photomultiplier tubes
\*\* Diode-pumped solid state

### Phenotyping of *CLV* mutants

For meristem measurements (area size, width and height) primary and secondary IFMs of wild type (*Col-0*) and mutant plants (*cle40-2, cle40-cr1-3, bam1-3, cle40-2;bam1-3, clv1-101*) were dissected at 6 WAG under LD conditions. For *clv3-9* and *clv1-101;bam1-3* only secondary IFMs were imaged and analyzed, due to the highly fasciated primary meristems. Optical sections of the Z-stacks were performed through the middle of the meristem starting in the centre of primordia P5 and ending in the centre of primordia P4. Based on the optical sections (XZ), meristem height and area size were measured as indicated in Fig6-SupplFig. 2. IFM sizes from Fig. *1*E are also used in Fig. *6*E for *Col-0*, *cle40-2* and *clv3-9* plants.

Same procedure was used to count the cells expressing *WUS* in different mutant backgrounds (Fig. 7A-E). Optical sections of IFMs at 5 WAG were performed from P4 to P5 and only nuclei within the meristem area were counted and plotted. For analyses of carpel numbers, the oldest 10 - 15 siliques per plant at 5 WAG were used. Each carpel was counted as one, independent of its size. N number depicts number of siliques. Leaf measurements were performed at 4 WAG and four leaves of each plant were measured and plotted. Data was obtained from at least 3 independent experiments.

### Data analysis

For visualization of images the open-source software ImageJ v 1.53c (Schneider et al., 2012) was used. All images were adjusted in “Brightness and Contrast”. IFMs in Fig. *7* were imaged with identical microscopy settings (except for *clv3-9* mutants) and were all changed in “Brightness and Contrast” with the same parameters to ensure comparability. *clv3-9* mutants were imaged with a higher laser power since meristems are highly fasciated. MIPs were created by using the “Z-Projection“ function and optical sections were performed with the “Reslice…” function resulting in the XZ view of the image. Meristem width, height and area size were measured with the “Straight line” for width and height and the “Polygon selection” for area size. The shape parameter σ was calculated by the quotient of height and width from each IFM. For L1 visualization the open-source software MorphoGraphX (https://www.mpipz.mpg.de/MorphoGraphX/) was used that was developed by Richard Smith. 2½ D images were created by following the steps in the MrophoGraphX manual (de Reuille et al., 2015). After both channels (PI and fluorophore signal) were projected to the created mesh, both images were merged using ImageJ v 1.53c.

For all statistical analyses, GraphPad Prism v8.0.0.224 was used. Statistical groups were assigned after calculating p-values by ANOVA and Turkey’s or Dunett’s multiple comparison test (differential grouping from p ≤ 0.01) as indicated under each figure. Same letters indicate no statistical differences. All plasmid maps and cloning strategies were created and planned using the software VectorNTI®.

## Acknowledgements

This study was funded by DFG through iGrad-Plant (IRTG 2466), CRC 1208 and CEPLAS (EXC 2048). We thank Cornelia Gieseler, Silke Winters, and Carin Theres for technical support and Yasuka L. Yamaguchi (Sawa lab), Anne Pfeiffer (Lohmann lab) and Rene Wink (Simon lab) for sharing Arabidopsis seeds. We also thank Vicky Howe for proof reading the manuscript, the Center for Advanced imaging (CAi) at HHU for microscopy support and Aleksandra Sapala for support with MorphoGraphX.

## Author contributions

J.S., G.D., Y.S. and R.S. designed and planned the experiments. J.S. performed experiments and data analysis, besides counting carpels (Fig1-SupplFig.3), which was performed by J. Schmidt. G.D., K.G.P. and J.S. generated stable Arabidopsis lines. Y.S. provided material. G.D., P.B. and J.S. performed the cloning. J.S. and R.S wrote the manuscript with input from all authors.

## Supplementary Figures

**Fig1-SupplFig.1:**
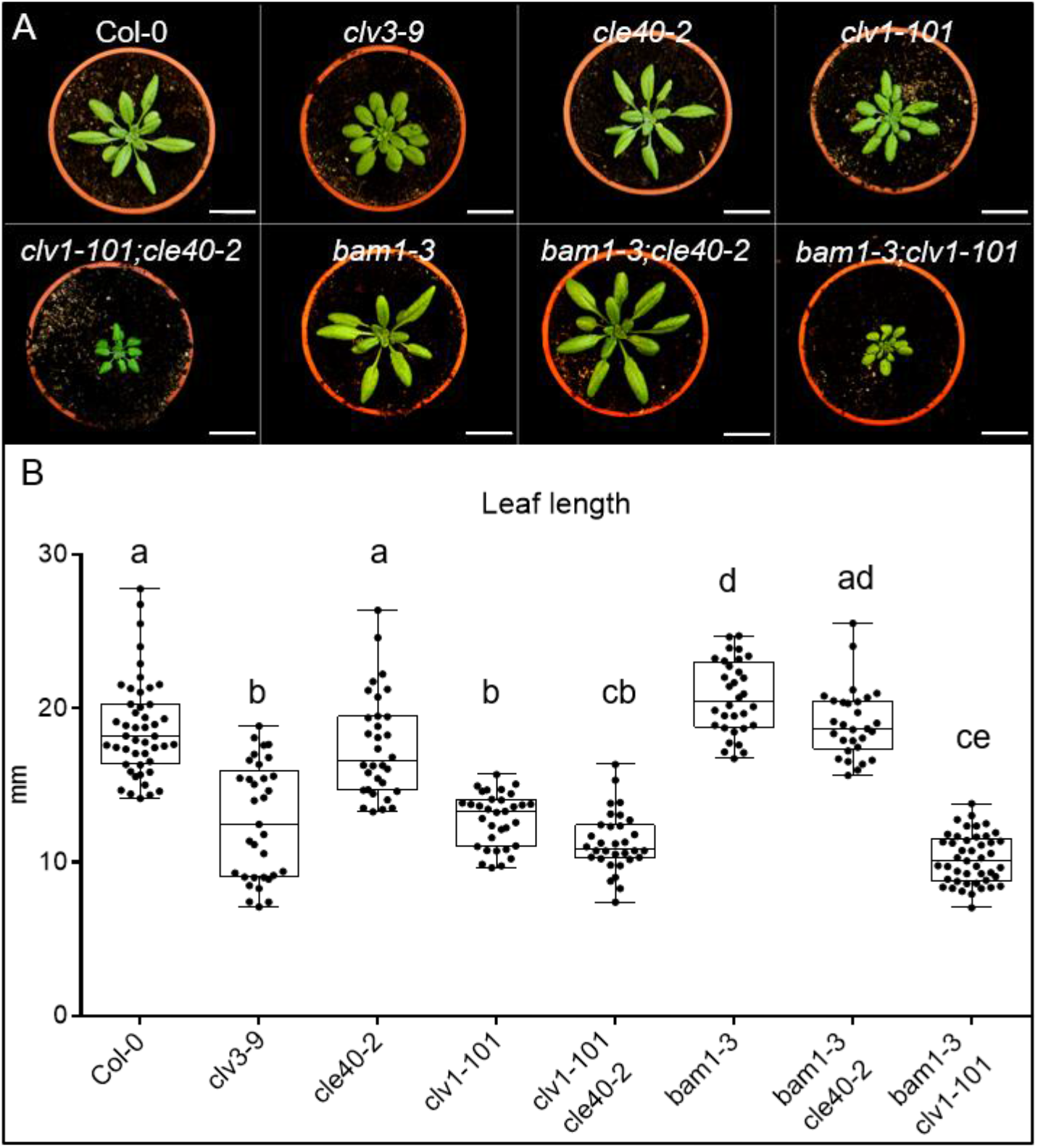
Mutants from the CLV pathway show differences in their leaf lengths. **(A)** Wild type (*Col-0*) and different single and double mutants (*clv3-9, cle40-2, clv1-101, clv1-101;cle40-2, bam1-3, bam1-3;cle40-2, bam1-3;clv1-101)* at 4 WAG. **(B)** Leaf lengths were measured and plotted. Wild type (*Col-0* N=47), *cle40-2* (N=32) and *bam1-3;cle40-2* (N=29) mutant plants do not show a significant difference in leaf length to each other. While *bam1-3* (N=32) mutants exhibit in average significantly longer leaves than wild type plants, the single mutants *clv3-9* (N=33) and *clv1-101* (N=33) and the double mutants *clv1-101;cle40-2* (N=32) and *bam1-3;clv1-101* (N=45) show significantly shorter leaves. Statistical groups were assigned after calculating p-values by ANOVA and Turkey’s multiple comparison test (differential grouping from p ≤ 0.01).

**Fig1-SupplFig.2:**
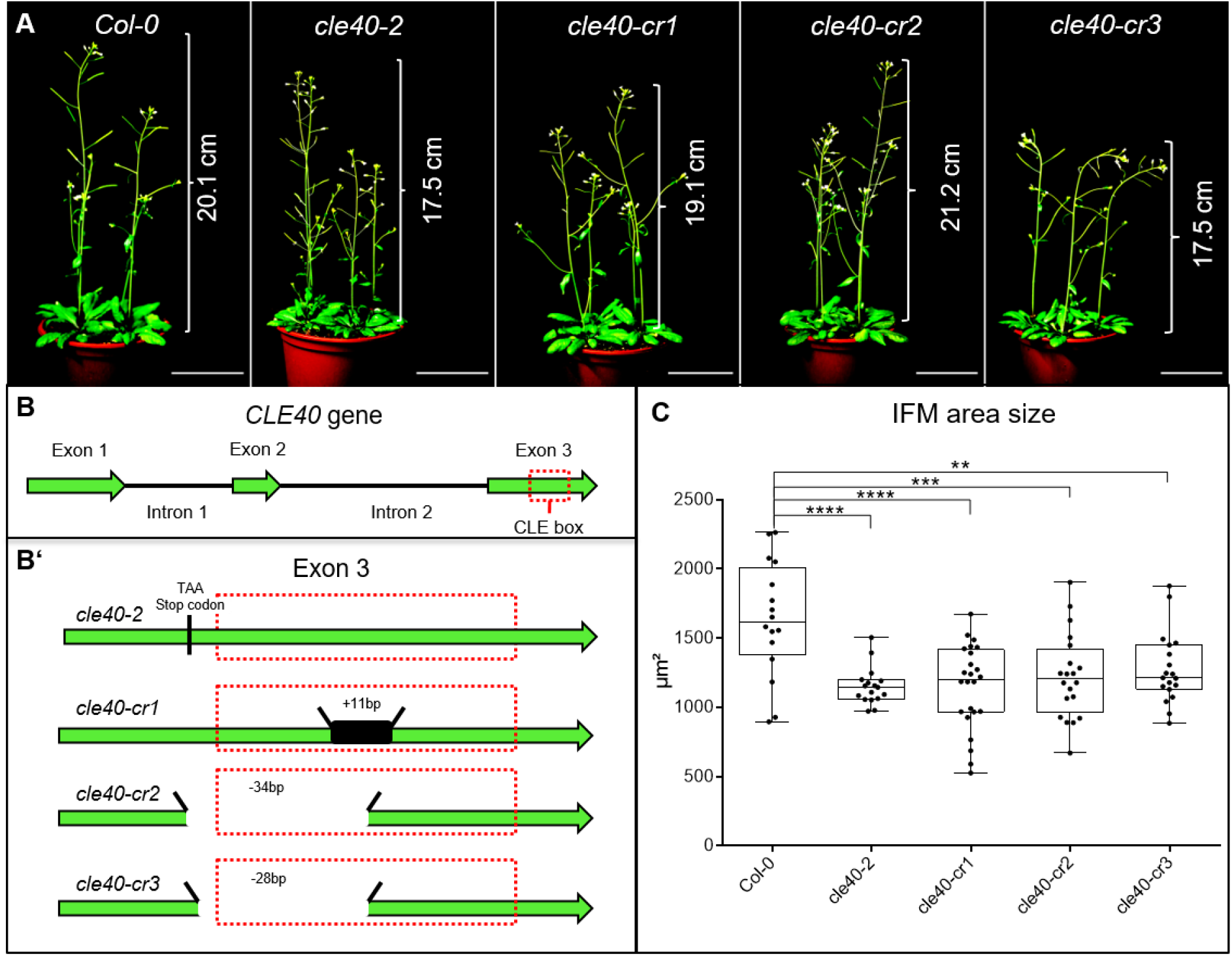
cle40 mutants have smaller meristems. **(A)** Wild type *A. thaliana* plants (*Col-0*) and *cle40* mutants (*cle40-2*, *cle40-cr1*, *cle40-cr2*, *cle40-cr3*) at 6 WAG. All plants show a similar height ranging from 17.5cm to 21.2cm and do not have an obvious plant phenotype. **(B)** Schematic representation of the *CLE40* gene, consisting of three exons (green arrows) and two introns. Exon 3 carries the crucial CLE box (dashed red line). **(B’)** Schematic representation of all four *cle40* mutations. All four lines have mutations in or before the CLE box domain in Exon 3. *cle40-2* mutants were created by transposon mutagenesis resulting in a stop codon in front of the CLE box (Stahl et al., 2009). *cle40-cr1*, *cle40-cr2* and *cle40-cr3* mutants were created using the CRISPR-Cas9 method (Yamaguchi et al., 2017). *cle40-cr1* has an 11bp insertion inside the CLE box domain while *cle40-cr2* and *cle40-cr3* have a deletion of -34bp and -28bp within the CLE box. **(C)** At 6 WAG, IFMs of wild type (*Col-0* N=16) and *cle40* mutant plants were dissected and the area of each meristem was imaged and measured. All four *cle40* mutants show significantly reduced IFM sizes compared to *Col-0* plants (*cle40-2* N=17, *cle40-cr1* N=24, *cle40-cr2* N=20, *cle40-cr3* N=19). Statistical stars were assigned after calculating p-values by ANOVA and Dunett’s multiple comparison test (differential grouping from p ≤ 0.01).

**Fig1-SupplFig.3:**
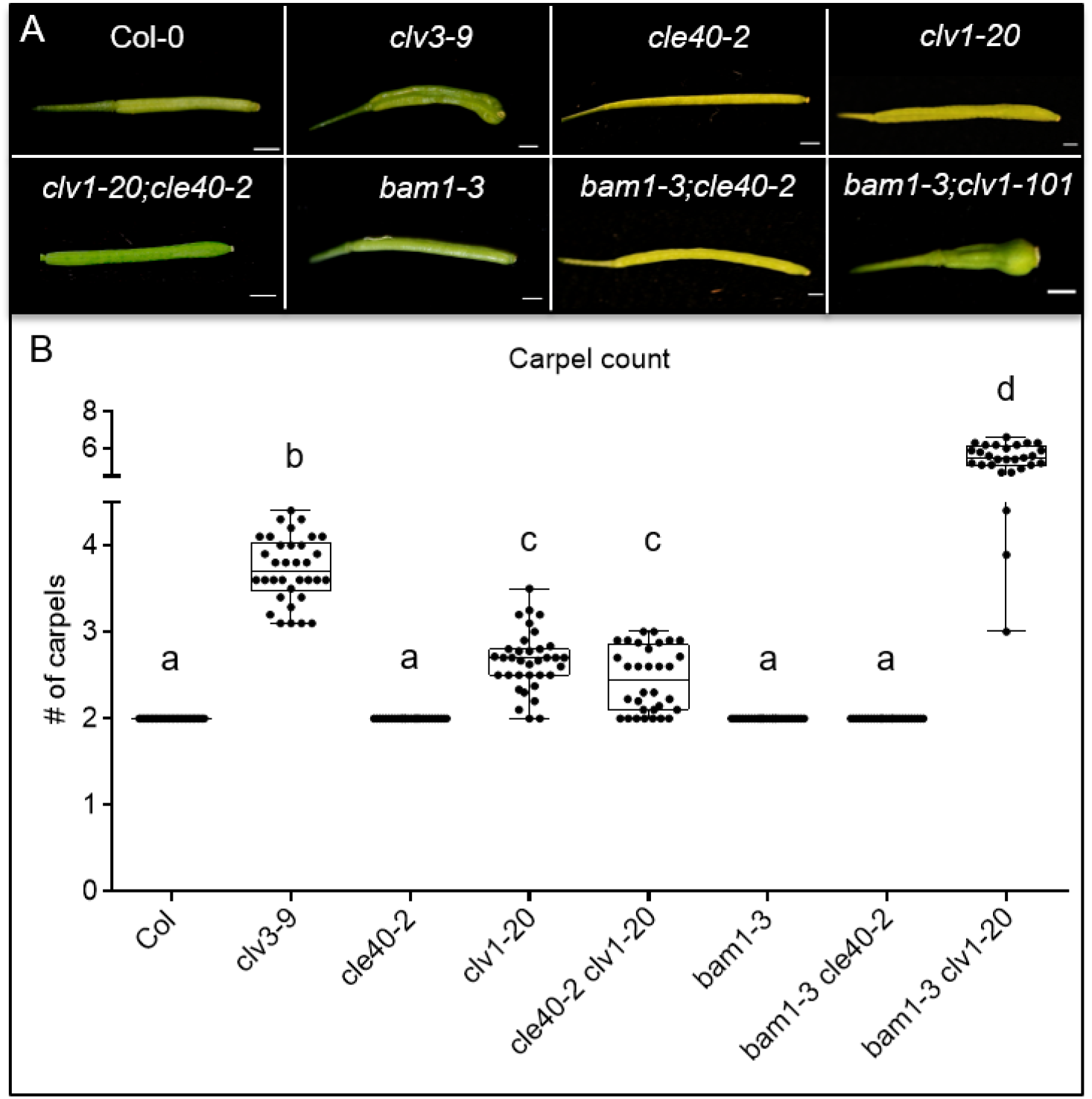
Siliques of various mutants differ in their carpel number. **(A)** Carpels of *Arabidopsis thaliana* plants at 6 WAG in wild type (*Col-0*) or different mutant backgrounds: *clv3-9, cle40-2, clv1-20, clv1-20;cle40-2, bam1-3, bam1-3;cle40-2, bam1-3;clv1-101*. **(B)** Carpel number was counted and plotted. Wild type (*Col-0* N=290), *cle40-2* (N=300)*, bam1-3* (N=300) and *bam1-3;cle40-2* (N=280) mutant plants always develop tow carpels, while *clv3-9* (N=340) plants exhibits 3 to 5 carpels and *clv1-20* (N=350) and *clv1-20;cle40-2* (N=320) mutants show in average 2 to 3 carpels. The double mutant *bam1-3;clv1-20* (N=280) develops 6 carpels in average. N number depicts number of siliques. Statistical groups were assigned after calculating p-values by ANOVA and Turkey’s multiple comparison test (differential grouping from p ≤ 0.01).

**Fig3-SupplFig.1:**
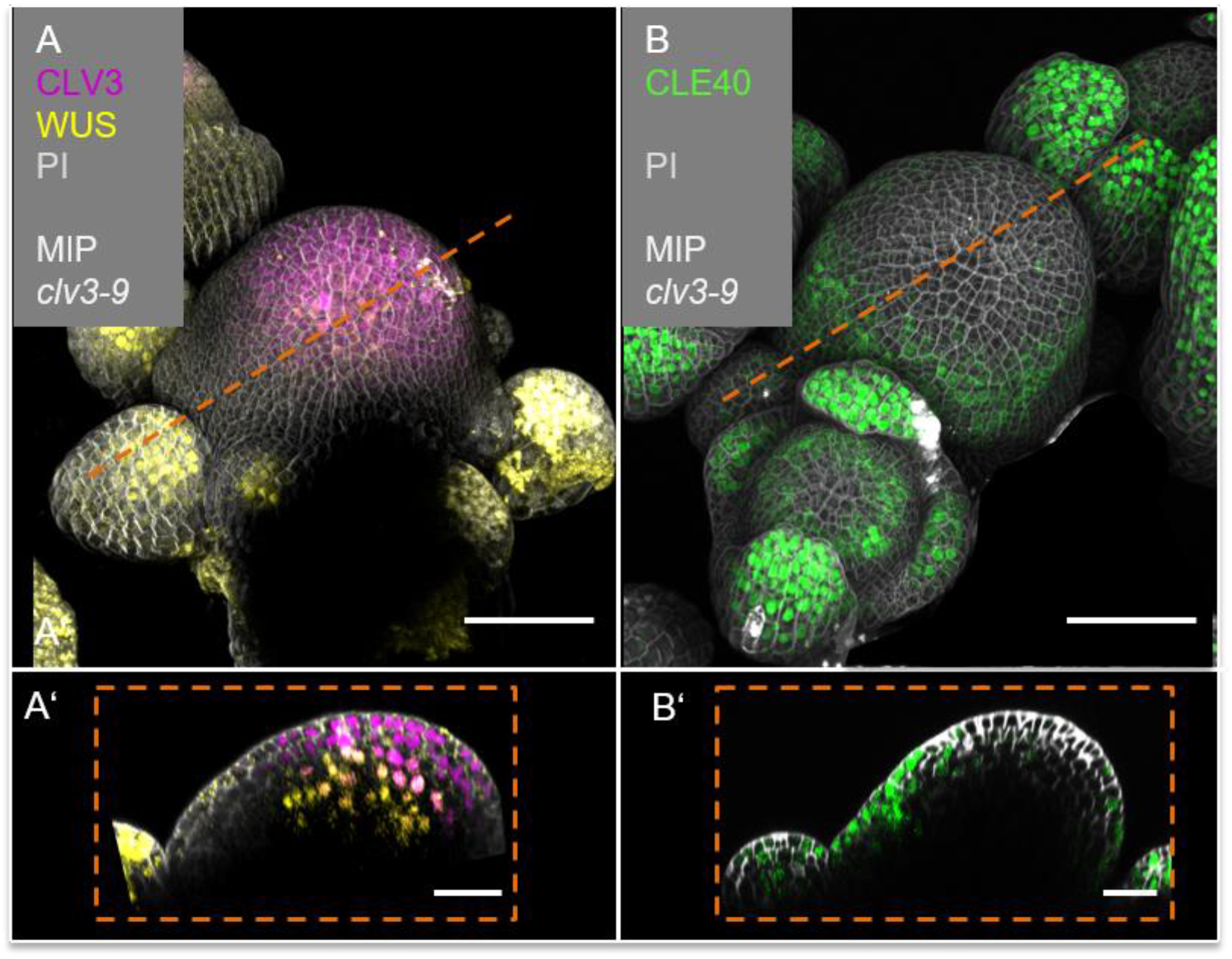
CLE40 expression is lacking in the CZ and OC. **(A)** MIP of *CLV3* and *WUS* expression (*CLV3:NLS-mCherry;WUS:NLS-GFP//clv3-9)* in a *clv3-9* mutant IFM (N=5). *CLV3* expression is detected at the tip of the meristem, while *WUS* expression is predominantly found in young primordia surrounding the meristem. **(A’)** Optical section through the IFM shows an extended expression domain of *CLV3* in the CZ and *WUS* expressing cells in OC of the IFM. **(B)** MIP of a *clv3-9* mutant IFM expressing *CLE40:Venus-H2B*. *CLE40* is expressed in the PZ of the IFM, in flower primordia and in mature sepals (N=6). **(B’)** Optical section through the IFM shows *CLE40* expression in the outer layers of the PZ while it is lacking in the CZ and OC, where *CLV3* and *WUS* are expressed. Dashed orange line indicates the planes of optical sections; Scale bars: 50µm (C, D), 10µm (C’, D’), MIP = maximum intensity projection, PI = propidium iodide

**Fig3-SupplFig.2:**
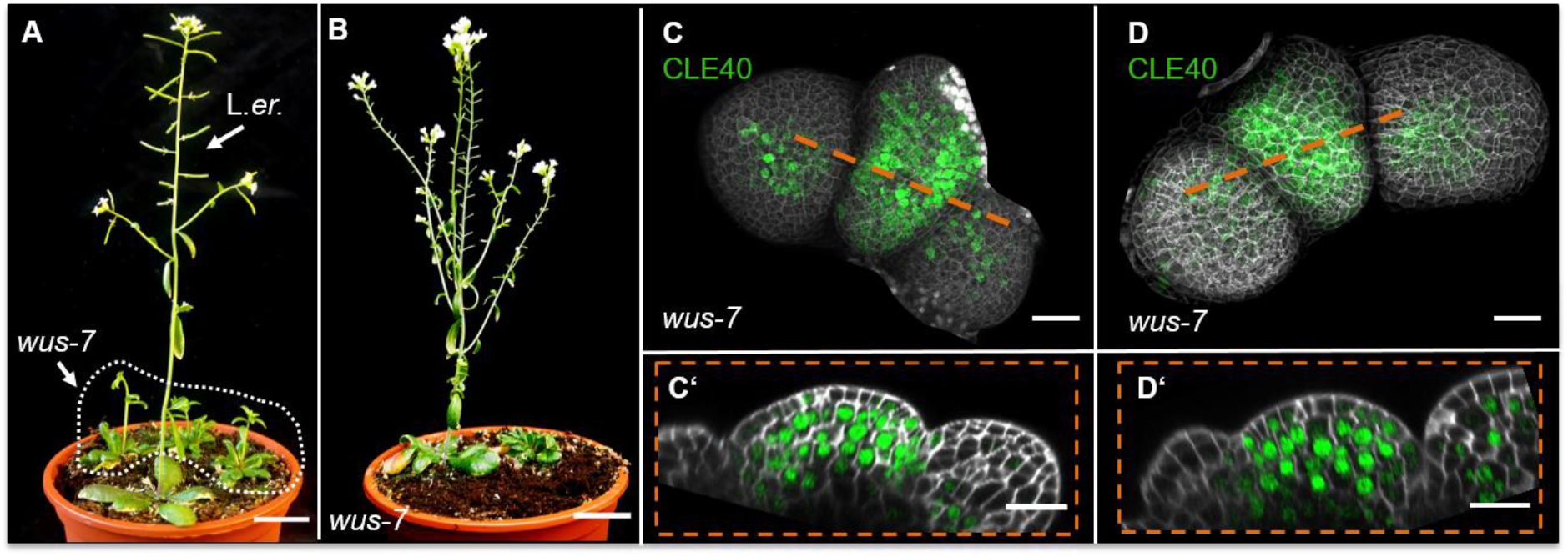
CLE40 expression is extended in *wus-7* mutants. **(A)** L.*er*. wild type plant at 5 WAG shows normal plant growth, while *wus-7* mutants at 5 WAG are delayed in their development (dashed white line). **(B)** *wus-7* mutant at 8 WAG. *wus-7* mutants develop IFMs but give rise to sterile flowers that lack inner organs. **(C and D)** MIP of *wus-7* IFMs at 5 WAG expressing *CLE40:Venus-H2B*. *CLE40* expression is detected through the entire meristem and in the centre of primordia (N=12). **(D’ and D’)** Optical sections through the meristem show *CLE40* expression in an extended pattern in the PZ and the OC. Dashed white line in B encloses homozygous *wus-7* mutants, dashed orange line indicates the planes of optical sections; Scale bars: 20mm (A, B), 20µm (C-D’)

**Fig6-SupplFig. 1.**
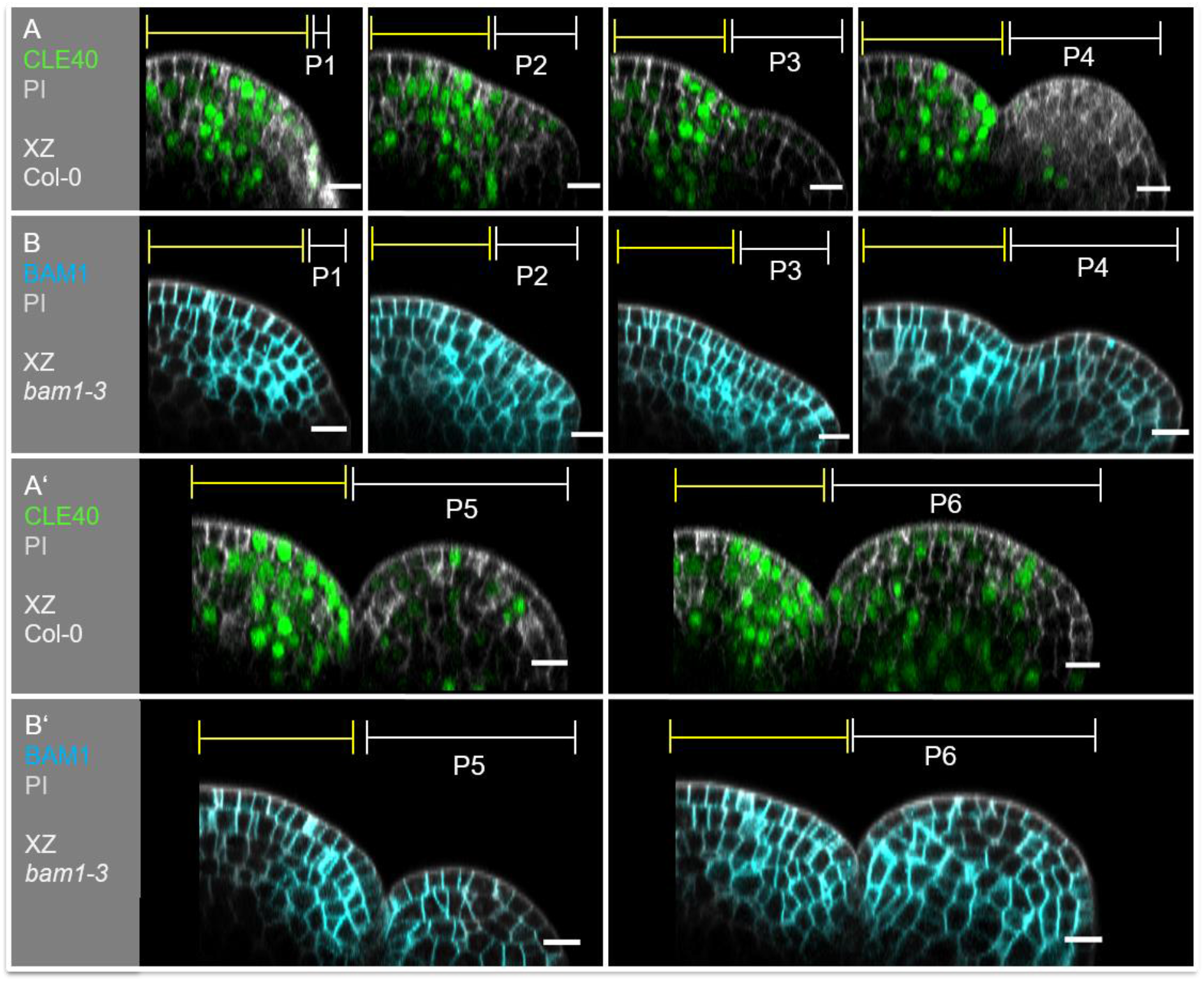
Expression patterns of *CLE40* and *BAM1* overlap in the IFM. Optical sections through an IFM and its developing primordia P1 to P6 expressing either **(A)** *CLE40* (*CLE40:Venus-H2B*) (N=23) or **(B)** *BAM1* (*BAM1:BAM1-GFP*) (N=15). In the IFM, *CLE40* and *BAM1* expression patterns overlap in the PZ, while both genes are lacking in the OC. No *CLE40* expression is detected in young primordia in P1 to P3. From P4 on a faint signal in CZ of the primordia express *CLE40*. Its expression expands in P5 and can be found in almost all cells of P6. *BAM1* is expressed ubiquitously in all primordia from P2 to P6. Yellow lines (P1 to P6) indicate the IFM region, white lines (P1 to P6) mark the primordium, Scale bar: 10µm, P = primordium

**Fig6-SupplFig. 2:**
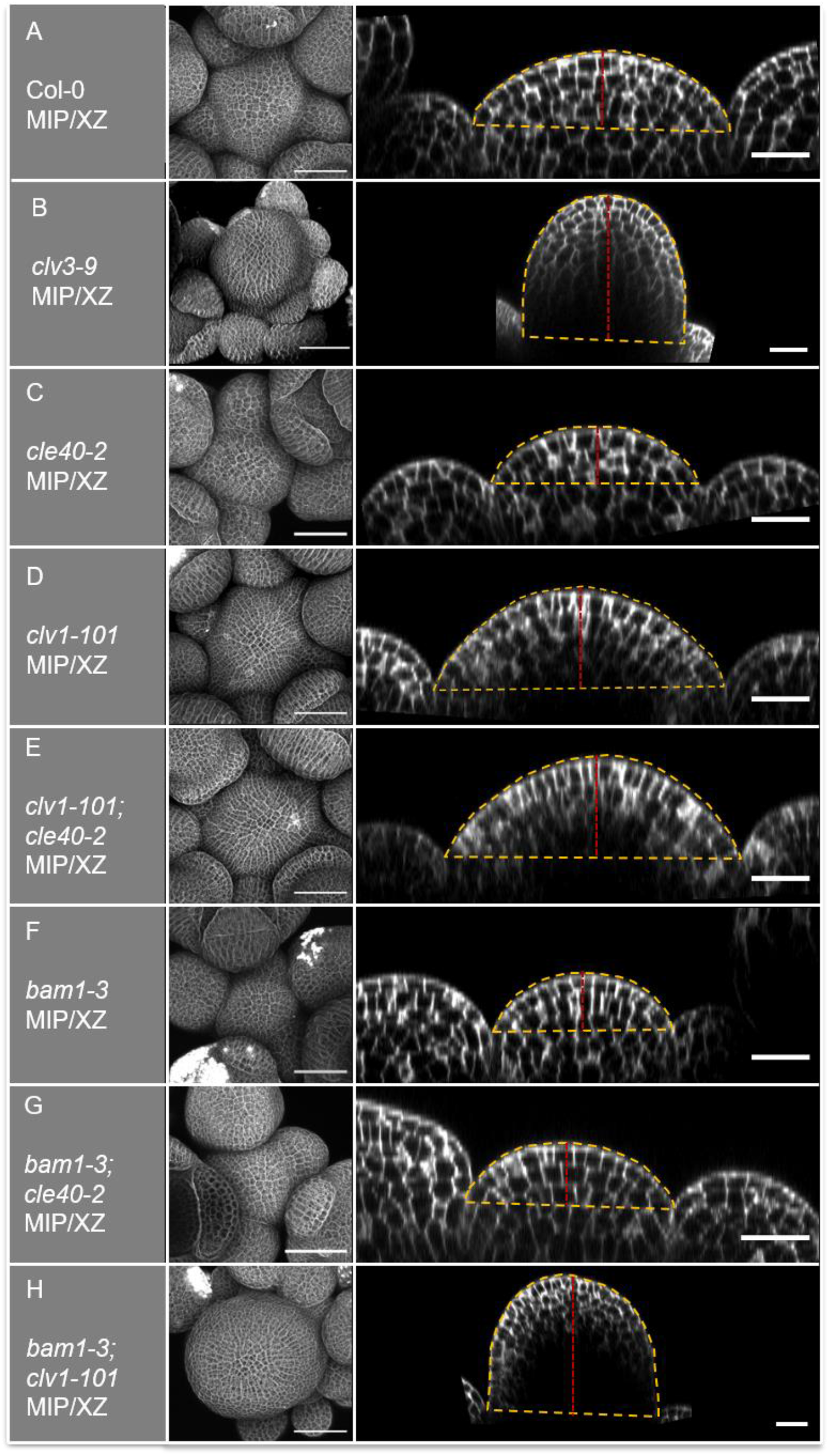
Meristem measurements in mutant backgrounds. IFMs of **(A)** *Col-0*, **(B)** *clv3-9***, (C)** *cle40-2*, **(D)** *clv1-101*, **(E)** *clv1-101;cle40-2*, **(F)** *bam1-3*, **(G)** *bam1-3,cle40-2* and **(H)** *bam1-3;clv1-101* were imaged after 6 WAG. Z-stacks were taken from the top of the IFMs with a confocal microscope. A MIP and an optical section from P4 to P5 was performed for each meristem. The yellow dashed line depicts the area of the meristem that was measured and the dashed red line indicates the height of the meristems. Scale bar: 20µm

**Fig7-SupplFig. 1:**
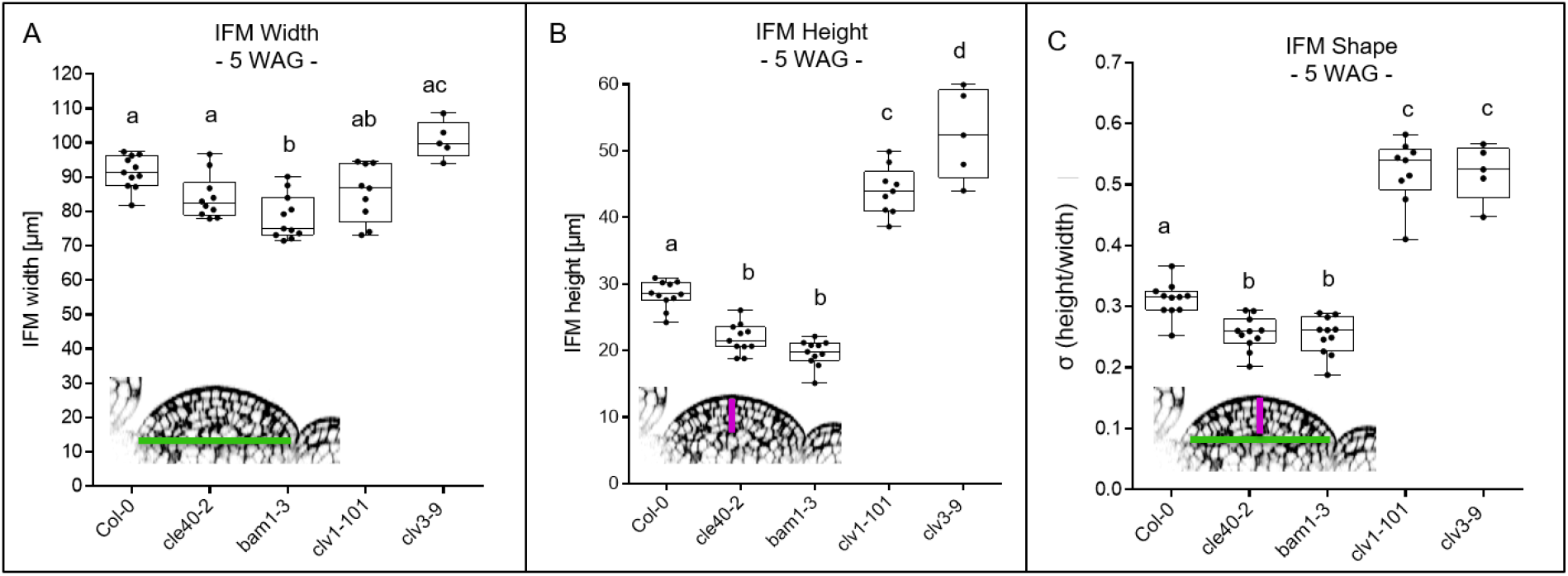
IFM height, width and shape after 5 WAG. **(A)** The width of *Col-0* (N=11)*, cle40-2* (N=10)*, bam1-3* (N=11), *clv1-101* (N=9) and *clv3-9* (N=5) mutants at 5 WAG does not significantly differ from each other. The average width lays between 85 to 100µm. Wild type plants are in average 90µm wide while *clv3-9* mutants depict the widest meristem average of 100µm. Only *bam1-3* mutants have with an average of 79µm a significantly smaller meristem wide compared to wild type plants. **(B)** The height of *cle40-2* (∼22µm) and *bam1-3* (∼23µm) mutants is significantly shorter after 5 compared to wild type plants (∼28µm). In contrast, *clv1-101* and *clv3-9* have significantly higher meristems than *Col-0, cle40-2* and *bam1-3* mutants. **(C)** The σ-value represents the shape of the meristem and is defined by the quotient of height and width. *cle40-2* and *bam1-3* mutants have a significantly smaller σ-value compared to wild type plants, resulting in flatter meristems. c*lv1-101* and *clv3-9* have with an average of 0.55 a significantly higher σ-value and thus have more dome-shaped meristems. Green line in the inset meristem in (A) indicates the width that was used for the quantifications in (A); magenta line in the inset meristem in (B) indicates the height that was used for the quantifications in (B); green and magenta line in the inset meristem in (C) indicates the width and height that was used for the quantifications in (C),

**Fig8-SupplFig. 1:**
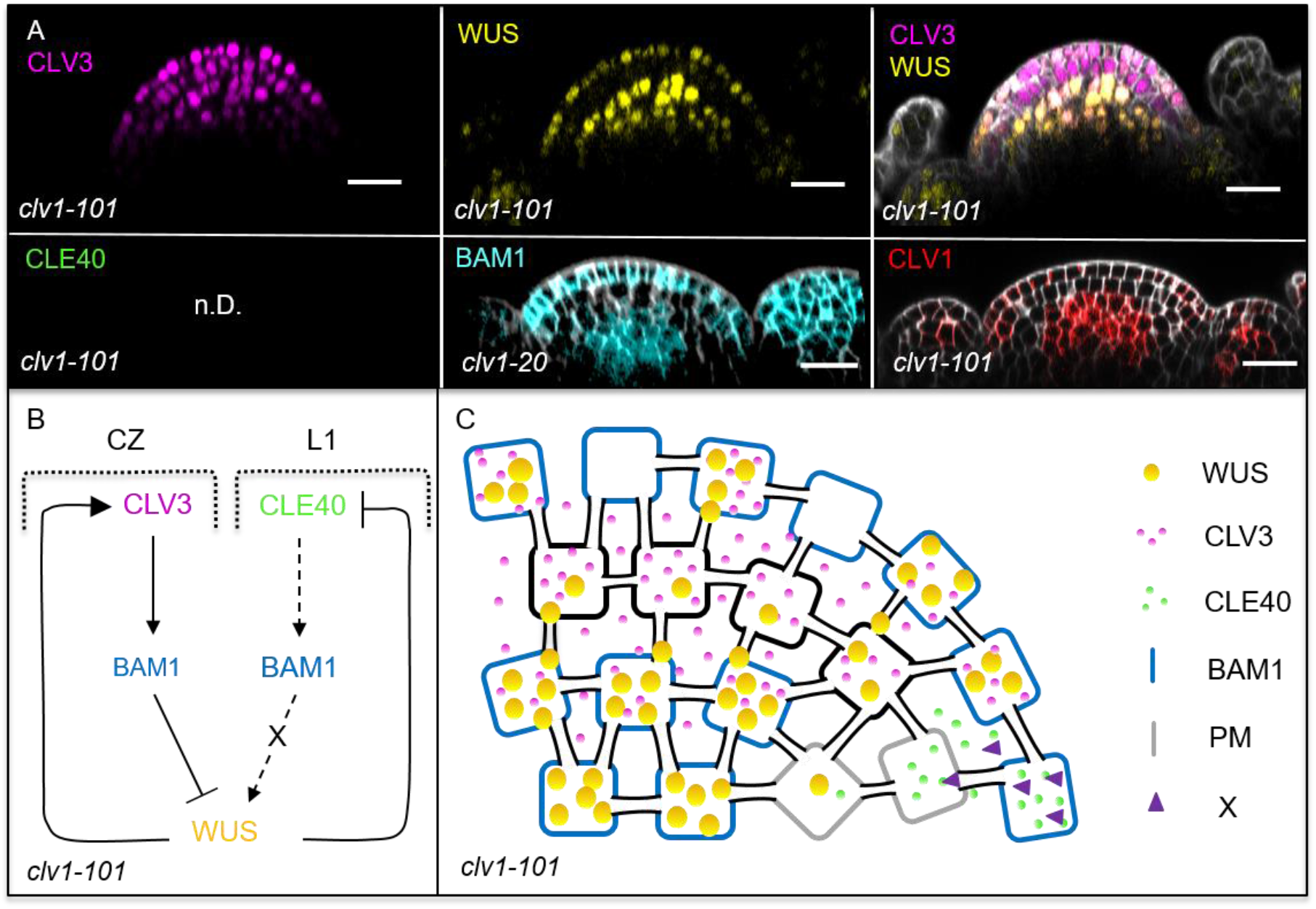
Schematic model of the two intertwined signaling pathways in a *clv1-101* mutant background. **(A)** Optical sections of through IFMs show the expression patterns of *CLV3* (N=8)*, WUS* (N=8)*, BAM1* (N=9) and *CLV1* (N=5) in a *clv1* mutant. Compared to wild type plants, the expression of *CLV3* and *WUS* is expanded and *WUS* is found in a patchy pattern in the L1. *BAM1* expression shifts to the CZ and is found in an elevated expression in the L1. **(B and C)** Schematic representation of two intertwined negative feedback loops in the IFM of a *clv1-101* mutant. The lack of CLV1 leads to a shift of *BAM1* expression to the OC and to an elevated expression in the L1. In the L3, BAM1 can partly substitute for CLV1 and thus CLV3 can act via BAM1 in order to repress WUS activity. The elevated expression of *BAM1* in the L1 overlaps in very few cells with *CLE40* expression in the periphery and leads to a weak activation of the downstream signal “X” that promotes WUS activity. Since, *WUS* expression is only partly repressed by the *CLV3-BAM1* signalling pathway, the WUS domain is extended and leads to an increase in stem cells (expanded *CLV3* expression). *WUS* is now also detected in the L1 of the meristem, together with *BAM1* expression. Scale bars: 20µm (A), CZ =central zone, L1 =layer 1

**Fig8-SupplFig. 2:**
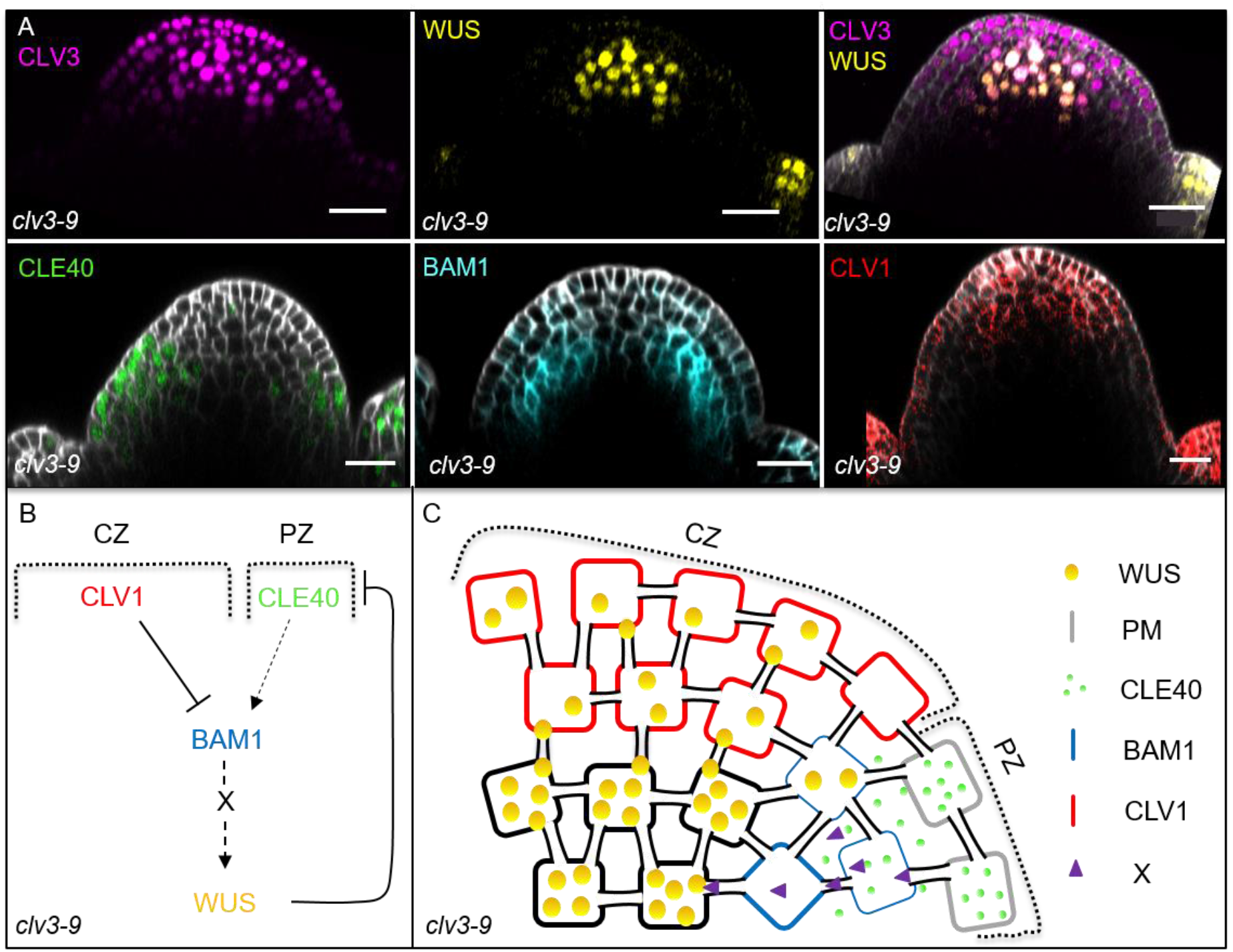
Schematic model of the intertwined signalling pathways in a *clv3-9* mutant background. **(A)** Optical sections of through IFMs show the expression patterns of *CLV3* (N=5)*, WUS* (N=5)*, CLE40* (N=6)*, BAM1* (N=5) and *CLV1* (N=5) in a *clv3-9* mutant. Compared to wild type plants, the meristem is highly increased in its size along the apical-basal axis and the expression of *CLV3* and *WUS* is expanded in the CZ and OC. *CLE40* expression is limited to the outer layers of the meristemś periphery and excluded from the CZ and OC, while *BAM1* expression shifts towards the inner layers of the PZ. *CLV1* expression is found at the tip and not in the centre of the fasciated meristem. **(B and C)** Schematic representation of two intertwined negative feedback loops in the IFM of a *clv3-9* mutant. The lack of CLV3 leads to a fasciated meristem with increased number of stem cells and thus an expanded CZ and a decreased PZ. Since no CLV3 peptide is available, CLV1 is not activated and expression of *CLV1* shifts from the OC to the tip of the CZ, where it represses *BAM1* expression. *BAM1* is expressed in the inner layers of the PZ, while *CLE40* expression is found in the outer layers of the PZ since it is repressed by the expanded WUS domain in the centre of the meristem. Thus only very few cells express both, *BAM1* and *CLE40* and hence, nearly no WUS promoting factor “X” is produced and the CLV3-CLV1 signaling pathway does not repress WUS activity. Scale bars: 20µm (A), CZ = central zone, PZ = peripheral zone

**Fig8-SupplFig. 3:**
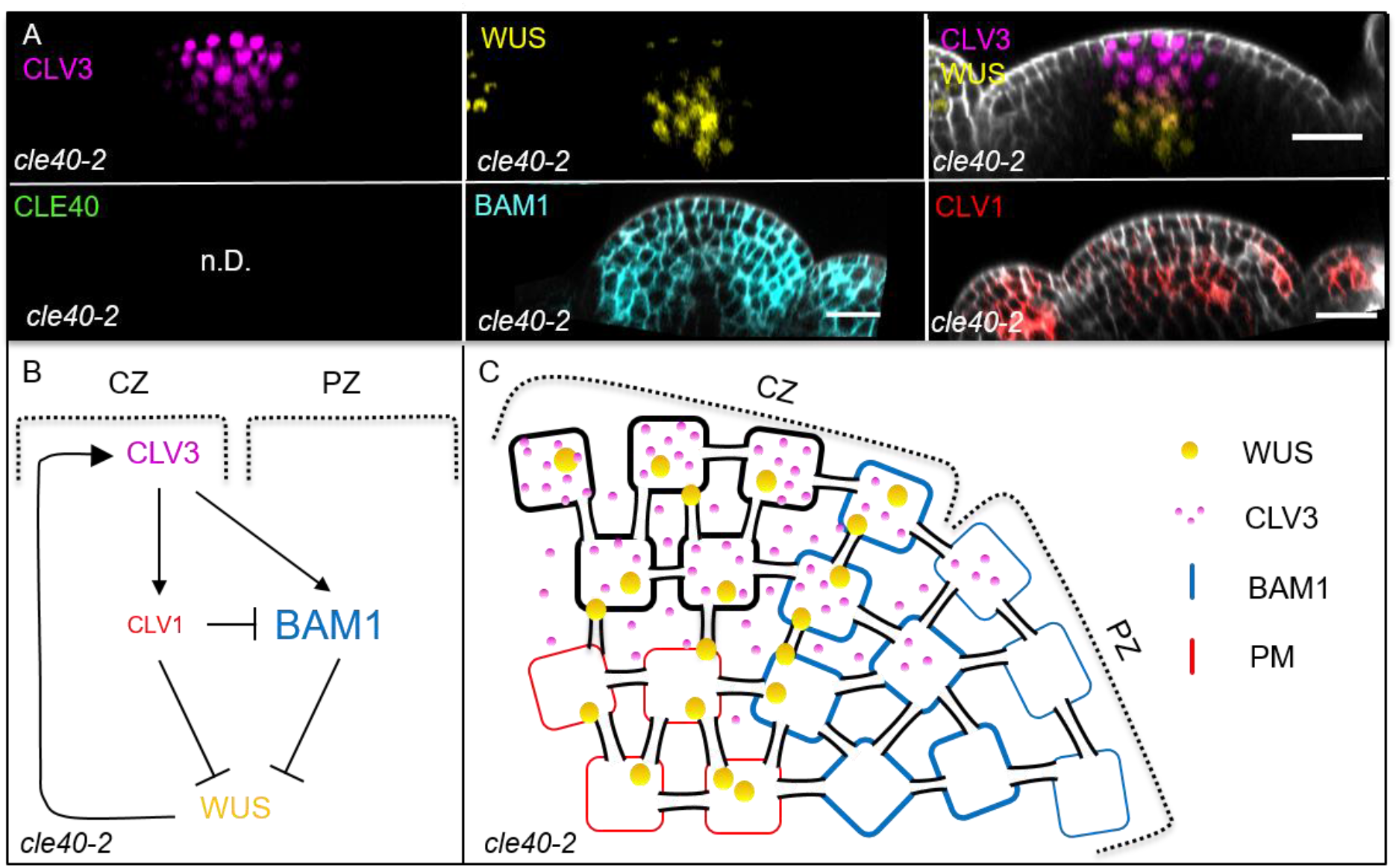
Schematic model of the intertwined signalling pathways in a *cle40-2* mutant background. **(A)** Optical sections of through IFMs show the expression patterns of *CLV3* (N=9)*, WUS* (N=9)*, BAM1* (N=7) and *CLV1* (N=9) in a *cle40-2* mutant. *CLV3* expression is similar to wild type plants, in the CZ. *WUS* expression is found in the OC, but in less cells than in *Col-0* plants. *BAM1* expression appears to be broader compared to wild type plants, while *CLV1* expression seems to be decreased in its intensity. **(B and C)** Schematic representation of two intertwined negative feedback loops in the IFM of a *cle40-2* mutant. In *cle40-2* mutants, *CLV1* expression seems to be decreased and leads to a broader *BAM1* expression compared to wild type plants. Since expression of *BAM1* is now also found in the CZ, CLV3 is able to bind CLV1 and BAM1 in the OC and CZ (respectively), leading to a double repression signalling cascade from the centre of the meristem. In the PZ, the downstream signaling cascade of BAM1 is not activated through CLE40 and thus the WUS promoting factor “X” is not being expressed and the WUS domain is confined to the centre of the OC. Scale bars: 20µm (A), CZ = central zone, PZ = peripheral zone

**Fig8-SupplFig. 4:**
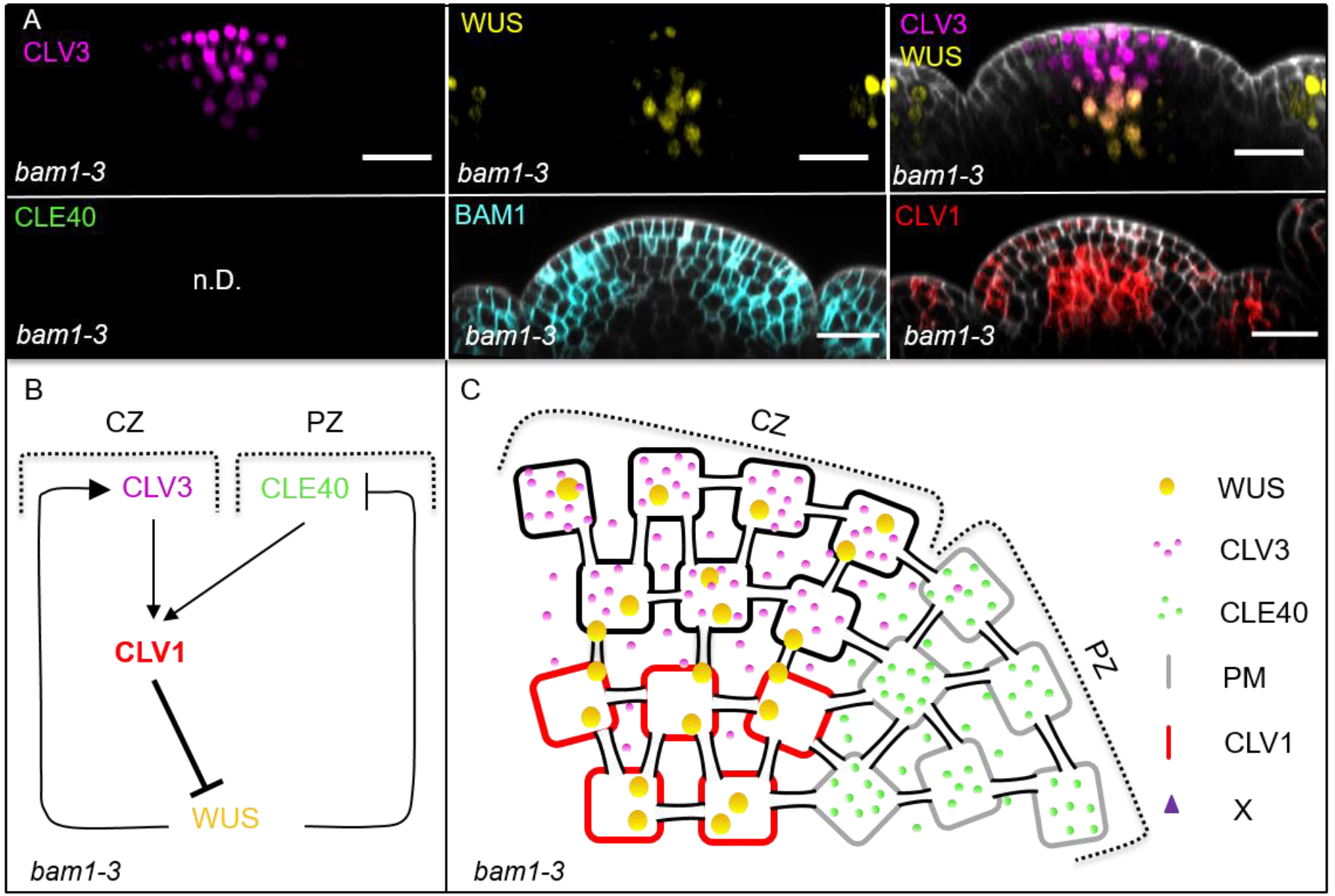
Schematic model of the intertwined signalling pathways in a *bam1-3* mutant background. **(A)** Optical sections of through IFMs show the expression patterns of *CLV3* (N=9)*, WUS* (N=9)*, BAM1* (N=15) and *CLV1* (N=7) in a *bam1-3* mutant. *CLV3* expression is similar to wild type plants, at the tip of the meristem in a cone shaped domain. *WUS* expression is found in the OC, but in less cells than in *Col-0* plants. *CLV1* expression seems to increased in its intensity compared to wild type plants. **(B and C)** Schematic representation of two intertwined negative feedback loops in the IFM of a *bam1-3* mutant. In *bam1-3* mutants, *CLV1* expression appears to be increased. Since BAM1 is lacking in the periphery, the *WUS* promoting diffusion factor “X” is not being produced and thus *WUS* expression is decreased and confined to the centre of the OC, similar to *cle40-2* plants. With the loss of BAM1, the main receptor for CLE40 is missing, and thus CLE40 peptide now might signal through CLV1 leading to a stronger repression of WUS from the centre of the meristem. Scale bars: 20µm (A), CZ = central zone, PZ = peripheral zone

